# A role for retro-splenial cortex in the task-related P3 network

**DOI:** 10.1101/2023.03.03.530970

**Authors:** Diptyajit Das, Marnie E. Shaw, Matti S. Hämäläinen, Andrew R. Dykstra, Laura Doll, Alexander Gutschalk

## Abstract

**Objective:** The P3 is an event-related response observed in relation to task-relevant sensory events. Despite its ubiquitous presence, the neural generators of the P3 are controversial and not well identified.

**Methods:** We compared source analysis of combined magneto- and electroencephalography (M/EEG) data with functional magnetic resonance imaging (fMRI) and simulation studies to better understand the sources of the P3 in an auditory oddball paradigm.

**Results:** Our results suggest that the dominant source of the classical, postero-central P3 lies in the retro-splenial cortex of the ventral cingulate gyrus. A second P3 source in the anterior insular cortex contributes little to the postero-central maximum. Multiple other sources in the auditory, somatosensory, and anterior midcingulate cortex are active in an overlapping time window but can be functionally dissociated based on their activation time courses.

**Conclusion:** The retro-splenial cortex is a dominant source of the parietal P3 maximum in EEG.

**Significance:** These results provide a new perspective for the interpretation of the extensive research based on the P3 response.

## 1. Introduction

Many tasks that we perform in response to sensory events recruit widespread cortical networks (Hugdahl et al., 2015; Kim, 2014) as detected by functional magnetic resonance imaging (fMRI). In electroencephalography (EEG), task-relevant stimuli ubiquitously evoke the prominent P3 (Sutton et al., 1965), also called P300, which has been explored by a large number of cognitive neuroscience studies including such diverse fields as consciousness (Sergent et al., 2005), mental disorders (Hamilton et al., 2020), or brain-computer interfaces (Chaudhary et al., 2016). Two variants of the P3 have been studied. The first is the earlier P3a, which is evoked by rare, salient events which are not assigned as target in an active task and is observed over more anterior sites in EEG. The second is the later P3b, which is only observed for task-relevant target events with an amplitude maximum over more posterior sites (Hillyard et al., 1971; Squires et al., 1975). The P3b is neither time-locked exactly to the stimulus nor to an optional motor response, but appears to represent a mapping between the two (Asanowicz et al., 2020; Verleger et al., 2014). The P3b has also been interpreted as a build-to-threshold process with a response increase until (briefly after) the motor response (O’Connell et al., 2012; Twomey et al., 2015). The latter is of interest for this study because of the characteristic response time course that is observed when the response is averaged to the motor response instead of to the target stimulus. More generally, there are numerous models of the potential psychological processes related to the P3, a summary of which is beyond the scope of this paper (Polich, 2007; Verleger, 2020). The focus of the present study is on the P3b, but the paradigm used will be expected to evoke some P3a as well, which is why we will refer to the response below simply as the P3.

Defining the functional role of the P3 in a neuroanatomically constrained model has been limited by ambiguous findings concerning its neural generators: Early intracranial EEG (iEEG) recordings in patients with epilepsy demonstrated P3-like responses in the hippocampus (Halgren et al., 1980), but subsequent studies in patients with lesions of medial temporal lobe structures demonstrated that the hippocampus is not the source of the P3 as measured by scalp EEG (Johnson, 1988; Onofrj et al., 1992). Further iEEG studies showed that P3-like responses can be observed by electrodes in many other brain areas (Halgren et al., 1995b, 1995a), and it was suggested that the neural generator of the P3 is distributed across multiple brain areas, including temporal, frontal, and parietal lobes, as well as the cingulate gyrus. This view was further supported by fMRI, which has been used to constrain source models of the P3 recorded in EEG (Bledowski et al., 2004; Li et al., 2020; Linden et al., 1999; C. Mulert et al., 2004). In agreement with other fMRI studies (Kim, 2014), these constrained source models suggested potential generators of the P3 in the pre-central sulcus, intra-parietal sulcus (IPS), supplementary motor area (SMA), midcingulate cortex (MCC), insular cortex, and temporo-parietal junction (TPJ). A potential role of the TPJ has been independently emphasized by studies in patients with structural brain lesions (Knight et al., 1989; Verleger et al., 1994; Yamaguchi and Knight, 1991). Source analysis of the P3 in magnetoencephalography (MEG) has typically suggested sources in the deep (Rogers et al., 1993; Tarkka et al., 1995) temporal lobes, but others reported that the P3 was not obtained reliably at all in MEG (Siedenberg et al., 1996). Currently, it is widely held that the P3 is generated by the same areas observed active during target detection of rare events in fMRI (Bledowski et al., 2004; Kim, 2014; Christoph Mulert et al., 2004), i.e., in TPJ, dorsal frontal and parietal cortex, and in the MCC and SMA. In other contexts, however, fMRI activity has been observed to co-vary with gamma-band activity rather than with evoked potentials in lower frequency bands (Logothetis et al., 2001; Niessing et al., 2005; Steinmann and Gutschalk, 2011) and a detailed investigation of how activity in defined anatomical areas would generate the spatial distribution of the P3 observed in EEG and MEG is still lacking.

The present study assessed the neural generators of the P3 by employing combined MEG and EEG (M/EEG) recordings and source analysis in a classical auditory oddball paradigm (Ritter et al., 1972), and directly compared such source analysis results to fMRI. Our results suggest a different source configuration for the P3 than summarized above, with one source lying in retro-splenial cortex (RSC), and another source lying in insular cortex. This P3 activity is paralleled by activity in multiple other areas, including auditory cortex (AC), primary somato-sensory cortex (S1), and anterior midcingulate cortex (aMCC), which can be dissociated from the P3 by their activation time courses. In the second part of the paper, we simulated the scalp EEG and sensor MEG based on circumscribed sources in these brain regions to evaluate (i) their contribution to the centro-parietal P3 that is typically evaluated in EEG, and (ii) to control for the interaction between remote source areas. Finally, we tested which of the simulated sources can explain the data at the P3 peak. Results suggest that the source in RSC explains more variance than other sources.

## 2. Methods

### 2.1. Participants

A total of fifteen healthy young adults (8 female, 7 male) with a mean age of 26.8 years (range 20 - 45) with no previous history of neurological or hearing disorder participated in this study. The data of three participants were excluded later from the analysis due to large measurement artifacts (n=2) and incomplete recording (n=1). The study was approved by the ethics committee of Heidelberg University, Germany and each volunteer provided written informed consent before participation.

### 2.2. Experimental design and procedure

Simultaneous MEG and EEG data were recorded while presenting a classical auditory oddball sequence consisting of frequent standard (1000 Hz) and rare (14%) deviant (900 Hz) tones, presented with an average inter-stimulus interval of 2 s, randomly jittered by ± 0.5 s. All tones were 75 ms long and gated on and off with a 10-ms-long hanning window. The stimuli were presented in three runs with a duration of 14 minutes each, comprising overall 180 deviants. The randomization was different for each run, but the same sequences were used for all listeners, presented with a short break between runs. Listeners were instructed to press a response button with their right index finger each time that they detected a deviant tone. In M/EEG, stimuli were presented diotically with ER-3 earphones (Etymotics Research, Elk Grove Village, IL, USA) via foam earpieces. The sound was presented at a level around 60 dB SPL, individually adjusted to be at a comfortable listening level.

The fMRI data were recorded from the same participants in a separate session; the order of fMRI and M/EEG was randomized such that about half of the participants started with M/EEG and half with fMRI. The stimuli were generated with the same parameters as described for the M/EEG above, but independently randomized. Three stimulus sequences with a duration of 13 minutes each were presented to all listeners with a short break between runs, comprising overall 162 deviants. As in M/EEG, listeners indicated deviant detection by pressing a response button. In fMRI, stimuli were presented diotically via MR-compatible S14 insert earphones (Sensimetrics Corporation, Gloucester, MA, USA), which attenuate the scanner noise by approximately 15-20dB. Sound level was individually adjusted to be at a comfortable listening level around 65 dB SPL; the level was chosen higher in fMRI compared to M/EEG to compensate for masking by the scanner noise. All sound stimuli were generated using PsychoPy software (www.psychopy.org) (Peirce, 2007).

#### 2.2.1 Data acquisition

M/EEG data were acquired at a sampling rate of 500 Hz, using a 160 Hz low-pass and no high-pass filter. Recordings were performed inside a four-layer magnetically shielded room (IMEDCO) via a Neuromag-122 whole-head system (MEGIN OY, Helsinki, Finland) equipped with 61 dual-channel planar first-order gradiometers. Participants head geometry (80 points on the head surface) and location of four head-position indicator coils were digitized together with the EEG electrode positions relative to a coordinate system spanned by the nasion and two pre-auricular points using a Polhemus Isotrack II digitizer (Colchester, VT, USA). The positions of the head-position-indicator coils inside the MEG dewar were obtained before the recordings. EEG data were recorded using an Easycap (Herrsching, Germany) M64 recording cap with a 64-channel 10%-system montage. The EEG was referenced to Pz, amplified with two 32-channel Neuroscan amplifiers, and analog-to-digital converted together with the MEG.

MRI data were acquired with a 3T Siemens Magnetom Trio scanner (Siemens Medical Systems, Erlangen, Germany) with a 32-channel head coil; fMRI data were acquired with an interleaved echo planar imaging (EPI) sequence (TR=2 sec, TE=30 ms, flip angle 80°) with 32 axial slices aligned along the anterior-posterior commissure line (3.99-mm slices, 3×3 mm^2^ in-plane resolution). Structural MRI images with the same field of view were obtained, including T1-weighted anatomical images (GR/MPRAGE, flip angle 9, echo time 2.63, repetition time 1570, resolution 1×1×1 mm^3^) and multi-echo fast low-angle shot (FLASH) sequences. These images were used for co-registration with subject-specific M/EEG and fMRI results to standard space and for creating realistic-shaped boundary-element head models. The three scanning runs lasted 13 minutes and 20 seconds each and there was a brief break between runs to restart the stimulation and communicate with the participant.

### 2.3 M/EEG data processing

Preprocessing of M/EEG data was performed using MNE software packages (http://martinos.org/mne)(Gramfort et al., 2013). For each recording (three runs per participant), first, a visual inspection of the raw M/EEG data was carried out to identify and mark time epochs as well as channels containing large artifacts or flat signals. Flat-signal channels were reconstructed using spherical spline interpolation. A separate denoising step was then performed only for the MEG data to reduce uncorrelated sensor noise and artifacts (i.e., flux jumps) using oversampled temporal projection (OTP) (Larson and Taulu, 2018). This technique allows suppression of sensor-space noise that is spatially uncorrelated with the data. The M/EEG data were then bandpass filtered (0.5-30 Hz) and the EEG was re-referenced to average reference. Eye blinks and cardiac artifacts were then removed from the data using MNE’s independent component analysis algorithm (Hyvärinen, 1999). Afterwards, the data were epoched from -100 to 1000 ms relative to stimulus onset, yielding two stimulus locked conditions: standard and deviant. A separate, response-locked epoching window was created, spanning -500 to 500 ms relative to the button press. Thus, three overall data conditions were constructed: standard, deviant (both stimulus locked), and response-locked. Deviants without subsequent button press (miss trials) and button presses after standards (false alarms) were excluded from the analysis. Next, artifact contaminated trials were repaired or excluded using the automatic data driven ’autoreject’ (Jas et al., 2017) algorithm implemented in MNE. The number of epochs for standards was reduced to the number of deviants using the ‘mintime’ function in MNE, to equalize the number of trials across conditions for the source analysis.

#### 2.3.1 The source space and gain matrix

To define an individual, cortically constrained source space, FreeSurfer (Dale et al., 1999; Fischl, 2012) was first used to reconstruct the cortical surface (white and pial) from the high-resolution T1-weighted scan (3D MPRAGE data) for each participant. Afterwards, 10242 sources per hemisphere were placed at the gray-white matter boundary to create a source space with 3.1 mm average distance between the two nearest sources. In this source space, each dipole source constitutes a cortical surface area of about 9.5 mm^2^ (average quotient across dipoles and participants). We then manually excluded sources in the corpus callosum, thalamus, and lateral wall of the ventricle (Glasser et al., 2016), because these areas do not comprise cortex and are thus unlikely to be sources of the M/EEG. This procedure reduced the number of sources to around 9300 per hemisphere.

High resolution inner-skull, outer-skull, and scalp surfaces created from FLASH images were used to model the electrical conductivity between each surface using a three-compartment boundary-element model (BEM). For BEM, 5120 triangles were used for creating the triangulated meshes with respective conductivities of the brain, skull, and skin assumed to be 0.3 S/m, 0.006 S/m, and 0.3 S/m. To define the locations of the EEG electrodes on the scalp and the configuration of the MEG sensors relative to the cortical surface, MNE-coordinate-system alignment tools (Gramfort et al., 2013) were used, where fiducial landmarks (two pre-auricular points and the nasion) are manually identified from the MRI-based rendering of the head surface (Besl and McKay, 1992). The tool calculates a transformation by minimizing the digitized scalp surface points with respect to the MRI-defined scalp.

#### 2.3.2 Inverse modeling and source analysis of M/EEG data

The field distribution y(t) of sensor/electrode space M/EEG data can be modeled as a linear combination of the source time courses x(t) and noise n(t):

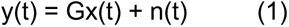

where, G is the forward gain matrix. To estimate the source current density on the cortical surface for each participant, an individual forward solution was computed (Gramfort et al., 2013; Hämäläinen and Sarvas, 1989; Uutela et al., 2001). The inverse estimation of active sources (x) is then performed by applying an inverse operator (G’) to the data by using the linear L2 minimum-norm estimator (MNE) such that:

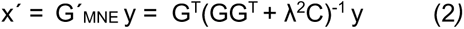

where, x’ is an estimation of the true sources x, C is the noise covariance matrix at the sensor/electrode space, and λ is the Tikhonov regularization parameter. In addition to that, a loose orientation constraint of 0.2 (0 = fixed orientation; 1 = free dipole orientation) was added to the model. This parameter has been empirically shown to improve source localization (Lin et al., 2006a). Afterwards, the source estimates were normalized to yield a dynamic statistical parametric map (dSPM) (Dale et al., 2000). The noise-covariance matrix was calculated from pre-stimulus baseline i.e., 100 ms preceding the stimuli by using an automated advanced regularization method called shrinkage technique (Engemann and Gramfort, 2015). Subsequently, noise-normalized source-space data from each participant were transformed onto a template brain atlas i.e., the FreeSurfer average brain (fsaverage) using a spherical registration method (Fischl et al., 1999). This registration was used to accurately align the dSPM results across individuals. The resulting maps across participants were then averaged per evoked condition to create a single grand-average dSPM solution.

#### 2.3.3 Regions of interest and source time courses for M/EEG data

A basic notion for the choice of regions of interest (ROIs) for the time-course analysis was the consideration that the polarity represented by the dSPMs can be interpreted by their relationship to the underlying sources if they were recorded with invasive EEG. Because the dSPMs are based on a dipole layer orthogonal to the cortical mantle, negative-going activity translates into surface negative activity with respect to the cortical surface, assuming that the activity would similarly be recorded as negative-going by invasive electrodes placed directly over the source on the cortical surface (Steinschneider et al., 1992). In the M/EEG view from a distance, limitations caused by cancelation between sources and the point-spread of the source analysis need to be additionally considered: Large sources of homogeneous polarity at the cortical convexity, which span one or more gyri and sulci, will be subject to considerable cancelation between sources on opposite sides of a gyrus/sulcus, such that radial sources are often dominating in non-invasive EEG in this case (Ahlfors et al., 2010). In the dSPM such activity typically maps more strongly to the radially oriented convexity of gyri and concavity in the depth of sulci. The other limitation of macroscopic M/EEG and distributed source analysis lies in the point-spread function of the technique (Hauk et al., 2022), meaning that a focal source (technically a dipole) is not mapped focal, but spreads around the source and across sulci and gyri that show a similar anatomical orientation. This spread to opposite banks of a sulcus or gyrus (or fissure) then shows opposite polarity to the source. A likely assumption for such a dSPM configuration is therefore that the source is limited to (or at least dominant at) one bank of a sulcus or gyrus with homogeneous polarity at a given latency. Critically, even such a restricted source area of homogeneous polarity in the dSPM is not directly related to the anatomical extent of an underlying source, but more to the orientation of the dipoles in the particular area of the distributed source model.

Accordingly, three constraints were used for ROI definition: (1) the source should be active in the grand-average dSPM maps, (2) the source should be active in the grand-average fMRI maps, and (3) the source should be limited to one side of a sulcus/gyrus or to the radial convexity of a gyrus and show homogeneous polarity in the dSPM grand-average maps. Note that the latter criterion is not intended to capture the extend of the proposed source (ROIs were typically smaller than the area active in fMRI), but to provide an approximation of the anatomical center of the expected neuro-electrical sources and their characteristic orientation that can be used to estimate source time courses. More details on the choice and definition of the ROIs used is provided in the results section.

ROIs were created on the FreeSurfer average brain and were then transformed to the individual brain anatomy using a surface based spherical morphing technique. As a consequence, not only the area but also the orientation of the associated sources was adapted to the individual anatomy. Source time courses were then calculated at the individual level as average of the noise-normalized activity in all dipoles comprised in a given ROI, using only the cortex normal orientation (i.e. not the loose constraint used for the dSPM maps). The mean-flip procedure of MNE python was used, which flips the orientation of dipoles whose orientations deviate more than 180 degrees from the average orientation of an ROI. Because of the polarity constraints used for the definition of ROIs, no dipole orientations were flipped with the exception of a few dipoles in S1, producing no meaningful difference between flipped and original orientations overall.

#### 2.3.4 Statistical tests and reproducibility

The statistical difference among ROI-based source-level time courses between standards and deviants was assessed through a cluster-based permutation test (Maris and Oostenveld, 2007) across participants. The statistical test is a non-parametric test that is designed to solve the multiple comparisons problem. In detail, first, an F statistic is computed at each participant-specific ROI-based source-space data sample (every 2 ms from -100ms to 1000ms relative to stimulus onset) from each data condition. A cluster threshold (p < 0.01) drawn from a standard F distribution was then applied at each sample, keeping only the statistically significant samples to form clusters whose values were higher than the applied threshold. These clusters were tested against a maximum cluster-level permutation distribution under the null hypothesis. The maximum cluster-level distribution was constructed by taking the maximum cluster-level statistics (i.e., sum of absolute test statistics) produced by the clusters under the permutations. A total of 5×10^5^ permutations were used and they were generated with random partitions of the data. Afterwards, cluster level p-values were estimated by computing the proportion that resulted in some larger cluster-level statistics than the actual one calculated from the maximum cluster-level permutation distribution. Significant cluster p-values were defined by correcting the p-values using a Bonferroni correction i.e., critical alpha level (α = 0.05) was set to (α*= α/n <0.0025, where n=20; 10 ROIs x both hemispheres).

#### 2.3.5 Cortical M/EEG source simulations

Bilateral, anatomically constrained sources were simulated individually for each participant’s cortical surface based on the ROIs used for the time-course visualization. For each anatomical ROI, all dipolar sources lying within were uniformly activated with a time-course of an arbitrarily-chosen half-sinusoidal wave with a base frequency of 5 Hz. Source currents for dipoles were then scaled such that the absolute value summed over all dipoles within an ROI amounted to 25 nAm at the peak. The polarity of the simulated waveform was adapted to the polarity observed in the source waveforms of the respective ROI (i.e., negative going with the exception of the insular cortex ROI). After the amplitude normalization, dipoles within left and right hemisphere ROIs were combined to yield a bilateral source configuration. To simulate realistic sensor level noise, the individual noise-covariance matrices based on the M/EEG data were used and scaled to a number of 200 averages. Afterwards, simulated sources were projected back to the electrode/sensor level by multiplying the forward matrix with the source data. Individual spatial maps for scalp-EEG, sensor-MEG, and cortical dSPM source estimates were computed at the peak of the simulated sources for each individual data set and then averaged across all 12 simulated data sets.

#### 2.3.6 Spread analysis of M/EEG source data

The point-spread function and cross-talk function (Hauk et al., 2011) were computed in order to characterize the leakage of current estimates between different ROIs for the experimental data. First, the dSPM-based resolution matrix was computed by multiplying the inverse operator to the forward gain matrix for each ROI. Afterwards, each ROI point-spread function and cross-talk function was extracted as the column and the row of that resolution matrix, respectively. This step was then repeated for each individual data set before averaging across participants. Finally, leakage of current estimates and the potential influence of one ROI to another were calculated using an absolute Pearson correlation test between ROI specific point-spread functions and cross-talk functions.

#### 2.3.7 Explaining M/EEG experimental data by simulated activity

Linear combinations of the simulated M/EEG patterns were used to explain the scalp/sensor-level M/EEG data at a single time point for the P3 and, as a control condition, for the N1 using the same source configurations in both cases. The M/EEG data were averaged across subjects at the individual peak latency of the P3 in EEG electrode Pz and at the individual peak latency of the N1 in EEG electrode Cz. The simulated sources (2.3.5) were averaged across participants in electrode/sensor space. The relative weight for each simulated (bilateral) ROI was determined with an ordinary least-squares (OLS) procedure to best explain the M/EEG data. This procedure was performed separately for P3 and N1; it can be written as:

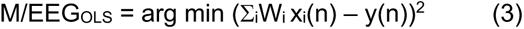

Where, W is the latent weighting vector for each (bilateral) ROI, x(n) is the ROI-based M/EEG simulation, and y(n) is the P3 (N1) data. For this procedure, the MEG and EEG data were normalized relative to the standard deviation and mean of the pre-stimulus baseline interval (i.e., MEG and EEG data were transformed to a z score), keeping the relative amplitudes within MEG and EEG data intact. To control for linear dependency, a multi-collinearity test was carried out by calculating the variance-inflation factor across the ROI-based M/EEG simulations. Variance-inflation-factor cutoff for linear independence was set to 5, i.e., all ROI combinations with a variance-inflation-factor value of less than 5 were considered linearly independent.

To quantify the quality of each model, the residual variance was calculated between the P3 (N1) data and the weighted combination of ROI-based simulations. The weights provided in the tables represent the weight W multiplied by 25 nAm, which was the strength of the summed simulated activity within each source. In this scaling, the weights provide a rough estimate of the source strength underlying the activity in the respective ROIs at the P3 (N1) peak latency. Note that this source strength does not equal the strength of a single dipolar source, since the variable geometry of ROIs cause different degrees of signal cancelation within each multi-dipole source.

### 2.4 fMRI data processing

For each participant, the functional volumes were mapped on the high-resolution anatomical surfaces using FreeSurfer. Surface-based fMRI data processing was then carried out using a standard FS-FAST routine (FreeSurfer’s functional analysis stream tool) (Fischl, 2012). First, preprocessing of the fMRI data was performed that includes the following sequence: template and brain-mask creation, followed by the registration of the functional data with FreeSurfer anatomical structure, motion correction, slice timing correction, intensity normalization of all voxels and time points, resampling of the data to the FreeSurfer average brain (fsaverage) atlas, and spatial smoothing of the data by a 5mm Full-Width/Half-Max (FWHM). Next, first level time-series analysis of the data was performed for each participant to remove nuisance variables (i.e., head motion) before computing p-values for a contrast between deviant and standard experimental conditions based on individual participant’s time courses with a canonical SPM hemodynamic response function. Later, a random-effects group analysis was performed across participants by using a Generalized Linear Model (GLM) implemented in FreeSurfer, followed by a multiple comparisons correction with the false-discovery rate (FDR, p<0.05) (Genovese et al., 2002) method. fMRI data processing steps were carried out for the left hemisphere and right hemisphere separately.

## 3. Results

A standard auditory oddball paradigm was used with the main goal of providing a high signal-to-noise ratio for source analysis. The paradigm comprised repeated, frequent 1000 Hz standard tones and rare 900 Hz deviants, which participants detected by button press. The average hit rate across subjects in M/EEG was 97±3% and mean reaction time 507±103ms (mean ± standard deviation). In fMRI the hit rate was 99±4% and the reaction time 490±150ms.

At the electrode and sensor levels (Fig. 1a), the two most prominent peaks of the event-related response are the earlier central negativity in both deviants and standards with a peak latency of 98 ms (mean ± 7 ms standard deviation) at electrode Cz (N1), and the prominent centro-posterior positivity evoked by deviants (targets) with a peak latency of 470 ms (mean ± 109 ms standard deviation) at electrode Pz (P3). While this P3 is readily evident as the biggest response in the EEG deviant waveforms, the N1 is more prominent in MEG. When averaging is aligned to the onset of the button press instead (Fig. 1b), the EEG shows a slow and steady increase up to 4 ms (mean±49 ms standard deviation) after the button press at Pz, whereas MEG activity shows a steeper increase right after the button press. Thereafter, EEG and MEG similarly show a slow and steady decrease. Maps of the EEG distribution at the individual maximum at Pz are highly similar when compared between the stimulus- and response-locked averaging. The MEG maps at the peak after the button-press show a dipolar pattern over the left central area, which appears to be somewhat different to the stimulus-locked averaging at the P3 latency as determined in EEG (Fig. 1c).

**Fig. 1.**
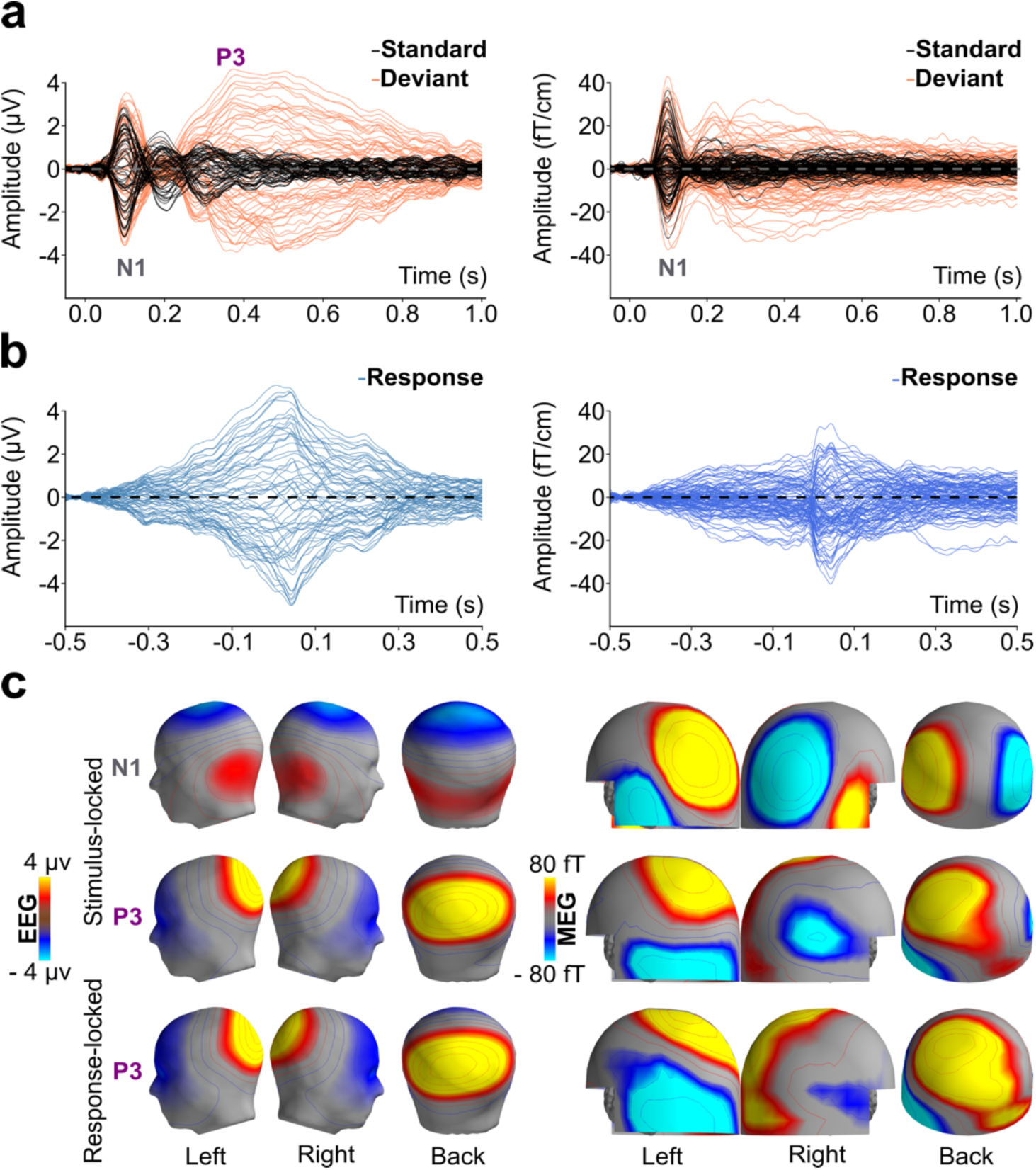
Grand-average evoked-response waveforms and maps. (a) Stimulus-locked EEG waveforms (left) and MEG waveforms (right) for standards (black) and deviants (coral). While the N1 is observed for standards and deviants alike, activity in the P3 time window from 300 – 600 ms is only observed for deviants. (b) Waveforms averaged to the button presses for detected deviants. While the EEG (left) is dominated by an increasing signal slightly beyond the button press, the MEG (right) shows a particularly strong, steeply rising response after the button press. (c) Grand average EEG maps (upper) and reconstructed MEG magnetometer maps (lower) are based on maps calculated at the individual peak latency of the N1 at electrode Cz (left), the P3 at electrode Pz (middle), and the response-locked average at electrode Pz (right).

### 3.1. Source analysis of the P3 in comparison to fMRI

To obtain reliable source models for the P3, the raw M/EEG data were first meticulously pre-processed to exclude, and model known artifact sources. Source analysis of the evoked response was obtained by calculating dSPM in an individual cortical source space, and then in an average across subjects by morphing the individual source estimates onto the Freesurfer average brain. The results of this procedure are shown in Fig. 2a. In the early N1 time range (T1: 75 – 125 ms), source activity is observed in AC in Heschl’s gyrus and planum temporale, with typical spread to adjacent and medial areas, including the inferior parietal lobes, superior temporal sulcus, medial temporal lobes, and posterior MCC (pMCC). For deviants, AC activity persists into the P3 time range (T2: 300 – 500 ms), but the activation pattern somewhat changes its distribution and extends more anteriorly towards the insular cortex then. Moreover, consistent activation is observed in the retro-splenial cortex (RSC) and in the posterior cingulate cortex (PCC). The RSC and PCC are opposite to each other, lying on the ventral and dorsal bank of the cingulate gyrus, respectively. Accordingly, the polarity of the mapped activity with respect to the cortical surface is positive in PCC and negative in RSC, suggesting that only one of the two areas is an active, biophysical generator of the P3. This activity continues into the later time window (T3:500 – 800 ms), in which additional activity is observed in anterior MCC (aMCC; subsumed to ACC in older anatomical nomenclatures). This aMCC activity is of opposite polarity with respect to the side of the cingulate gyrus when compared to PCC/RSC activity; i.e., aMCC activity is negative in the dorsal and positive in the ventral bank of the cingulate gyrus. Qualitatively similar source analysis results were also obtained by application of other widely used source estimation methods (Fig. S1).

**Fig. 2.**
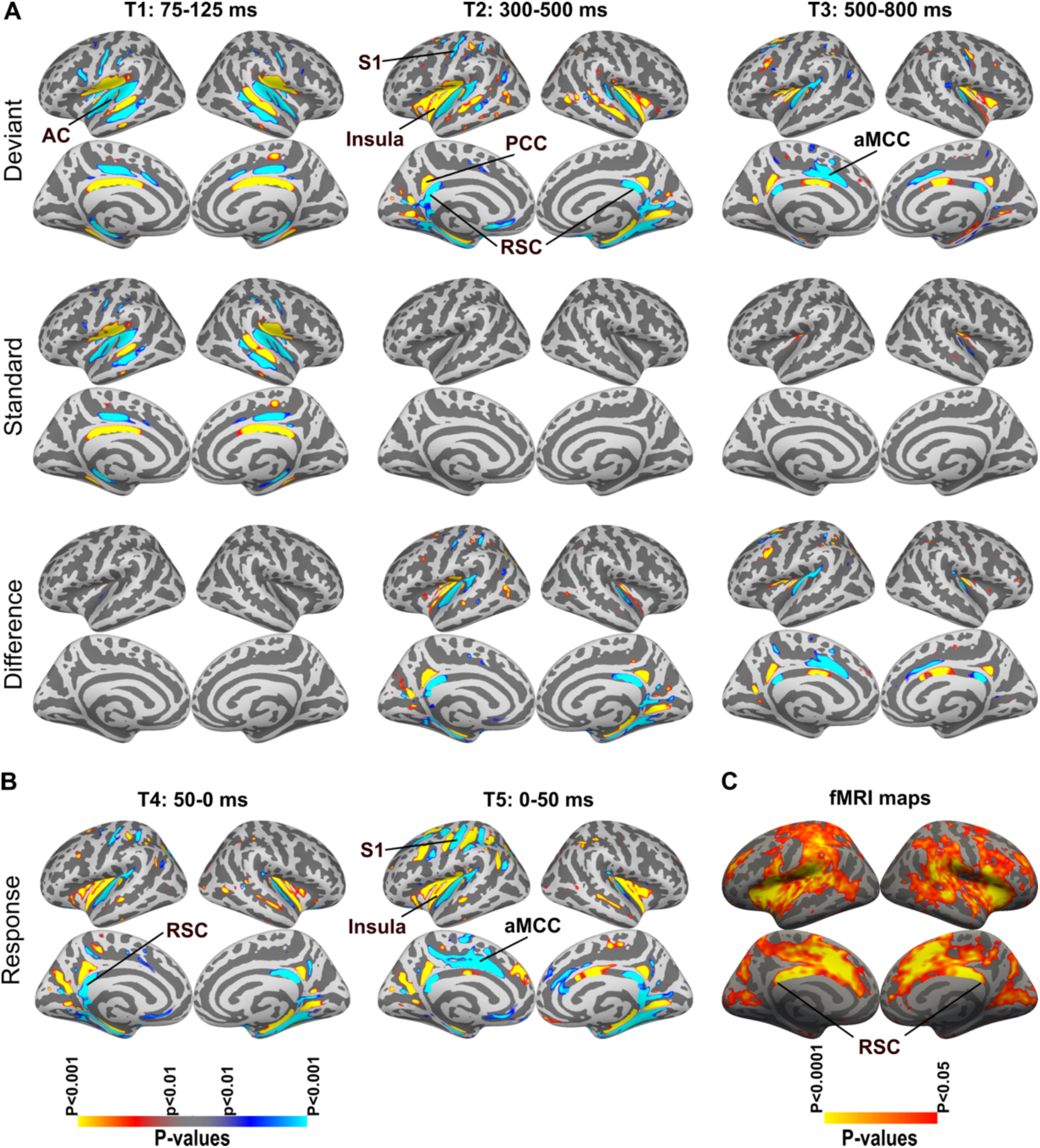
Cortical M/EEG and fMRI activation maps. (a) Dynamic statistical parametric maps (dSPM) based on the combined M/EEG for deviants (upper), standards (middle), and the contrast deviants – standards (lower) in three different time windows (n=12; p<0.01). The early 75 – 125 ms time window (T1) is focused on the N1, the middle 300 – 500 ms time window (T2) on the P3, and the late 500 – 800 ms time window (T3) on the late frontal negativity. Because the dSPMs are based on a fixed-effects statistic, the number of standard trials was reduced to the number of deviant trials for this analysis. (b) dSPMs for the response-locked average in the time window 50 ms before (T4) and 50 ms after (T5) the button press. (c) fMRI maps for the contrast deviants – standards (n=12; p<0.05, corrected for multiple comparison with the false-discovery-rate method), based on a random-effects statistic.

We also evaluated the activity averaged with respect to the button presses indicating correct target detection (Fig. 2b). The time windows evaluated were chosen here as 50 ms before and after the button press, to map motor and sensory activity related to the button presses. The maps for these response-locked averages overall show similar activation patterns as the stimulus-locked averages in RSC and insular cortex. In the 50 ms before the button press, no activity was observed in motor cortex or SMA. In the 50 ms after the button press, the activity around left S1 is very prominent and much stronger than in the stimulus locked averages. Moreover, activity in left aMCC was quite prominent after the button presses, extending somewhat into SMA and pre SMA.

A similar auditory oddball paradigm was employed in an fMRI experiment, to directly compare M/EEG and fMRI maps. The fMRI results for the deviant-minus-standard contrast (p<0.05, Fig. 2c) was generally consistent with previous reports of extensive brain activation for oddball, or generally target detection, with activity in frontal, parietal, and temporal lobes, as well as extensive activation in midline structures around MCC, pre-SMA, and SMA. When comparing the difference maps for M/EEG (Fig. 2a) and fMRI (Fig. 2c), it becomes evident that only part of the sites identified by fMRI also show significant source activity in M/EEG, including AC, MCC, S1, and insular cortex. Strong fMRI activity is also observed in RSC, but not in PCC. Based on this intermodal comparison, it would therefore appear that the M/EEG activity is also more likely generated in RSC. Some activity is also observed in the primary visual cortex (V1) and the parieto-occipital sulcus (POS) in fMRI and less consistent in M/EEG (POS activity only on the left). When the fMRI activity is thresholded more conservatively (Fig. S2), it appears that the most robust foci of activity are the RSC, aMCC/pre-SMA, insular cortex, auditory cortex, and TPJ. This pattern is quite similar to the M/EEG source analysis, with the exception of TPJ, where no significant activity was observed in the source analysis.

### 3.2. Source time courses of M/EEG

In order to explore the temporal characteristics of the prominent M/EEG sources in more detail, we calculated source-level time courses for regions that were active in both, M/EEG and fMRI. ROIs were restricted to areas of homogeneous polarity in the dSPM. The AC ROI was restricted to an area in dorsal Heschl’s gyrus, after the exploration of other regions, including planum temporale, had produced similar waveforms. An ROI in S1 was restricted to an area of the somatosensory hand area. The insular cortex ROI was chosen in the area where the dSPM maps showed positive polarity, and was restricted to the lower part where the deviant minus standard contrast showed significant M/EEG activity. Finally, two midline ROIs were defined: one in RSC on the ventral part of the cingulate cortex. This was motivated by the strong fMRI activity in RSC, while similar time courses were generally obtained with an ROI in PCC. An ROI in aMCC was defined in the dorsal side of the cingulate cortex, based on stronger fMRI activity on this side. The fMRI activation also extended into pre-SMA, but as pre-SMA activity was not clearly present in the dSPM maps, no additional ROI was added here.

As is demonstrated in the associated source time courses (Fig. 3), these ROIs segregate a number of distinct neural processes by their timing. The stimulus-locked averages (Fig. 3a) show the typical N1 waveform in AC. Subsequent to the N1, there is a sustained field that is significant in the deviant-minus-standard comparison on the left. Typical broad-peaked, P3-like time courses are observed in RSC and insular cortex, and with longer latency in aMCC. Note, however that the location of the RSC and aMCC ROIs are on opposite sides of the cingulate cortex and thus show opposite orientation towards the scalp, such that only the RSC can produce a positive-going field at Pz based on the negative-going late activity seen in the waveforms.

**Fig. 3.**
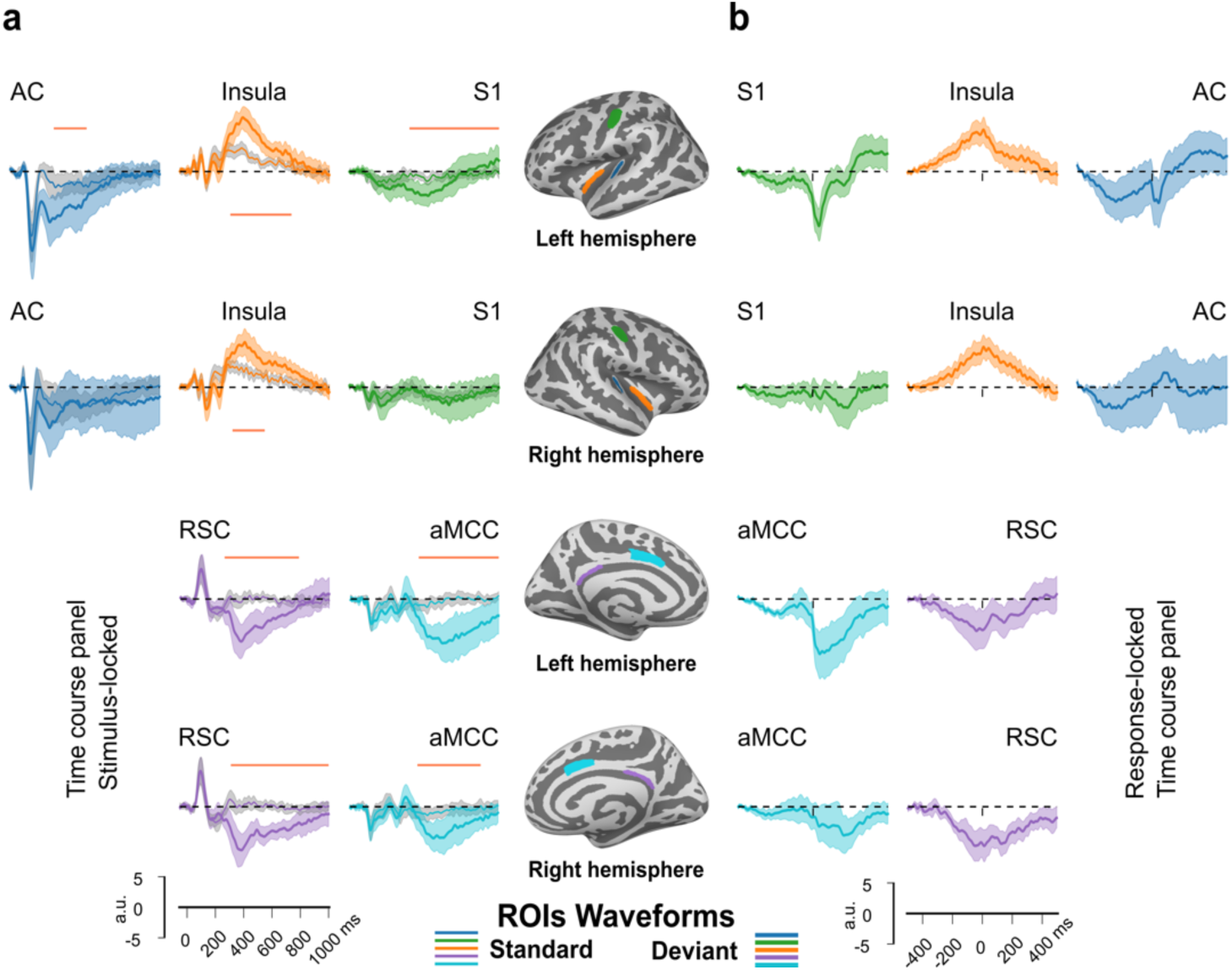
Region-of-interest (ROI) based source waveforms (average across participants, n=12; shaded area indicates 95% confidence interval). Source waveforms are based on dynamic statistical parametric maps (dSPM), calculated for the ROIs shown in the middle column with the same color code as the waveforms. The ROIs include auditory cortex (AC), anterior insular cortex (insula), primary somatosensory cortex (S1), retro-splenial cortex (RSC), and anterior midcingulate cortex (aMCC). (a) stimulus-locked source time courses, averaged relative to tone onset. Typical P3 source waveforms are observed in RSC (purple) and insula (orange). The coral color bar indicates the time interval in which the deviant and standard responses are significantly different from each other (cluster-based permutation test, see methods for details). (b) response-locked source time courses shown in similar configuration.

In the response-locked waveforms (Fig. 3b), the steadily increasing amplitude up to the button response, which dominates the scalp EEG waveforms (Fig. 1b), is observed at the source level in RSC and insular cortex. In contrast, activity in S1 shows a prominent transient wave that peaks approximately 40 ms after the button press, most likely representing tactile and proprioceptive somatosensory feedback related to the button press. A small transient after the button press is also observed in left AC; the latency of this wave coincides with the activity in S1, suggesting that it rather represents spread from or coactivation with S1 rather than auditory evoked activity related to the button press. Finally, activity in aMCC increases rapidly around the button press and persists for more than 300 ms thereafter.

To provide a more direct comparison of M/EEG and fMRI, we used activation clusters from the fMRI maps to extract stimulus-locked time courses from the M/EEG data. The results (Fig. S3) confirm a peak around 400 ms in RSC, and less prominent in the insular cortex. Time courses in aMCC/pre-SMA show a later peak around 600 ms. The TPJ time courses, in contrast, were dominated by activity that resembled the N1 and sustained field from AC, rather than the P3, irrespective of whether the orientation of the ROI was constrained or not. Results in the regions not explored in Fig. 3 were mixed: some P3-like patterns were observed in V1 and in the (left) POS, in particular when the orientation was not constrained. In frontal ROIs, consistent P3 peaks were not observed.

### 3.3. Comparison with simulated source data

Next, to evaluate the relationship between neural sources, spread of the source estimates, and scalp/sensor distributions, we computed (i) the source analysis and (ii) the scalp/sensor distribution for simulated data that would be generated by activity at the different ROIs based on the individual anatomy of the study participants. Each simulated source had a summed source current of 25 nAm (see methods section for details). As expected, these simulations show considerable spread from a focal source to neighboring sulci, for example to the inferior parietal lobe and to the superior temporal sulcus in the case of activity in the primary AC (Fig. 4a). Spread from AC is also observed in the medial temporal lobe around the hippocampal gyrus and in the pMCC, matching the activity pattern observed at the peak of the N1 in the original data (Fig. 2a).

**Fig. 4.**
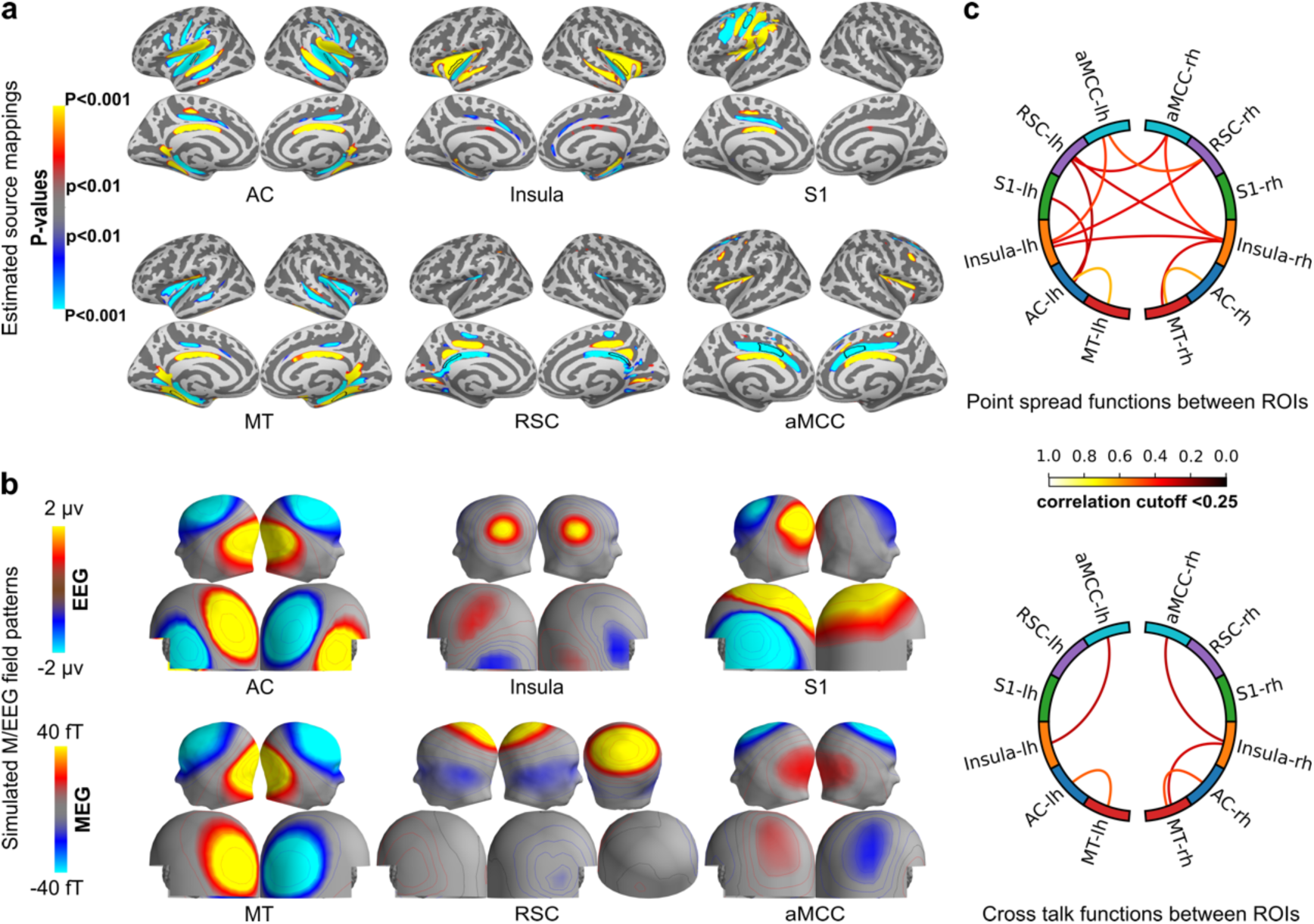
Simulated M/EEG and analysis of spread. (a, b) The data represent an average of n=12 individual simulations, based on bilateral, individually morphed regions of interest (ROIs) in auditory cortex (AC), Insula, medial temporal cortex (MT), retro-splenial cortex (RSC), and anterior midcingulate cortex (aMCC). The simulation of the primary somatosensory cortex (S1) is based on the left hemisphere ROI, only. Source polarity with respect to the cortical surface was chosen to match the pattern observed in the N1 (AC, MT) or P3 (insula, S1, RSC, aMCC) time window as shown in Fig. 2. (a) dynamic statistical parametric map (dSPM) of the simulated M/EEG data (p<0.01). (b) Maps of the scalp EEG (upper) and virtual magnetometer maps of the MEG based on the grand average of the simulated data, same scaling for all conditions. (c) Point-spread analysis (upper circle) and cross-talk analysis (lower circle) for the same ROIs used in (a) and (b). Each analysis is visualized using a circular graph with an absolute arbitrary correlation cutoff value of 0.25. The exact values are summarized in Fig. S4 (point spread) and Fig. S5 (cross-talk).

Some spread is also observed from AC to insular cortex and vice versa. The positive-going activity in insular cortex is not explained by spread from Heschl’s gyrus, however, suggesting that this activity observed in the P3 time window is really generated in insular cortex. To quantify the interaction between the evaluated brain regions in the dSPM source analysis, we calculated the point-spread function and cross-talk function between ROIs (Fig. 4c; Fig. S4 and Fig. S5). A strong interaction between sources within one hemisphere is observed between (1) Heschl’s gyrus and hippocampal gyrus, (2) insular cortex and aMCC, and (3) insular cortex and hippocampal gyrus. Strong spread is also observed between RSC and contralateral aMCC. Spread between left S1 and AC is also confirmed by this analysis, which may explain the S1-like waveform for response-looked waveforms in AC (Fig. 3a). In contrast, there was comparatively little spread between AC and insular cortex ROIs.

The RSC source produces a symmetric, posterior EEG scalp distribution that matches well with main aspects of the typical P3 observed in our data (Fig. 1c). In comparison to EEG, the simulated MEG activity for an RSC source is relatively weak. However, the simulated MEG map does not match well with the measured MEG map at the P3 peak (Fig. 1c). Moreover, the P3-peak maps showed higher reproducibility in EEG at the individual level, and much more variability in MEG (Supplementary Figures 5 and 6). These data indicate that the explanation of the EEG and in particular the MEG data requires multiple sources, for which MEG and EEG supposedly have different sensitivity.

### 3.4. Explaining the P3 with simulated M/EEG data

We therefore explored how the combined M/EEG data at the P3 peak can be explained by a combination of the sources that were used for the time-course analysis. To quantify the relative contribution of potential sources, the simulations based on the ROIs used for the time-course analysis were fitted with a least-squares procedure to the P3 data at its peak. The results show that a very good explanation of EEG and moderate explanation of MEG data can be achieved with this procedure (Fig. 5), leaving 6% residual variance in EEG and 62.5% residual variance in MEG (Table 1). Among all five sources, the RSC was scaled to the highest amplitude. When one of the sources was systematically omitted from the model, a massive increase of residual variance in EEG was only observed with the RSC omitted. Note that omitting RSC also led to the strongest increase of residual variance in MEG, supporting that the relatively weak contribution of the RSC is still relevant for MEG. All other sources only caused weak increment of the residual variance in EEG when omitted. This was also the case for the S1 source, the omittance of which increased the residual variance by only 0.4% in EEG, but by 19% in MEG, demonstrating that this source is almost as important for the MEG maps as the RSC at the P3 peak.

**Fig. 5.**
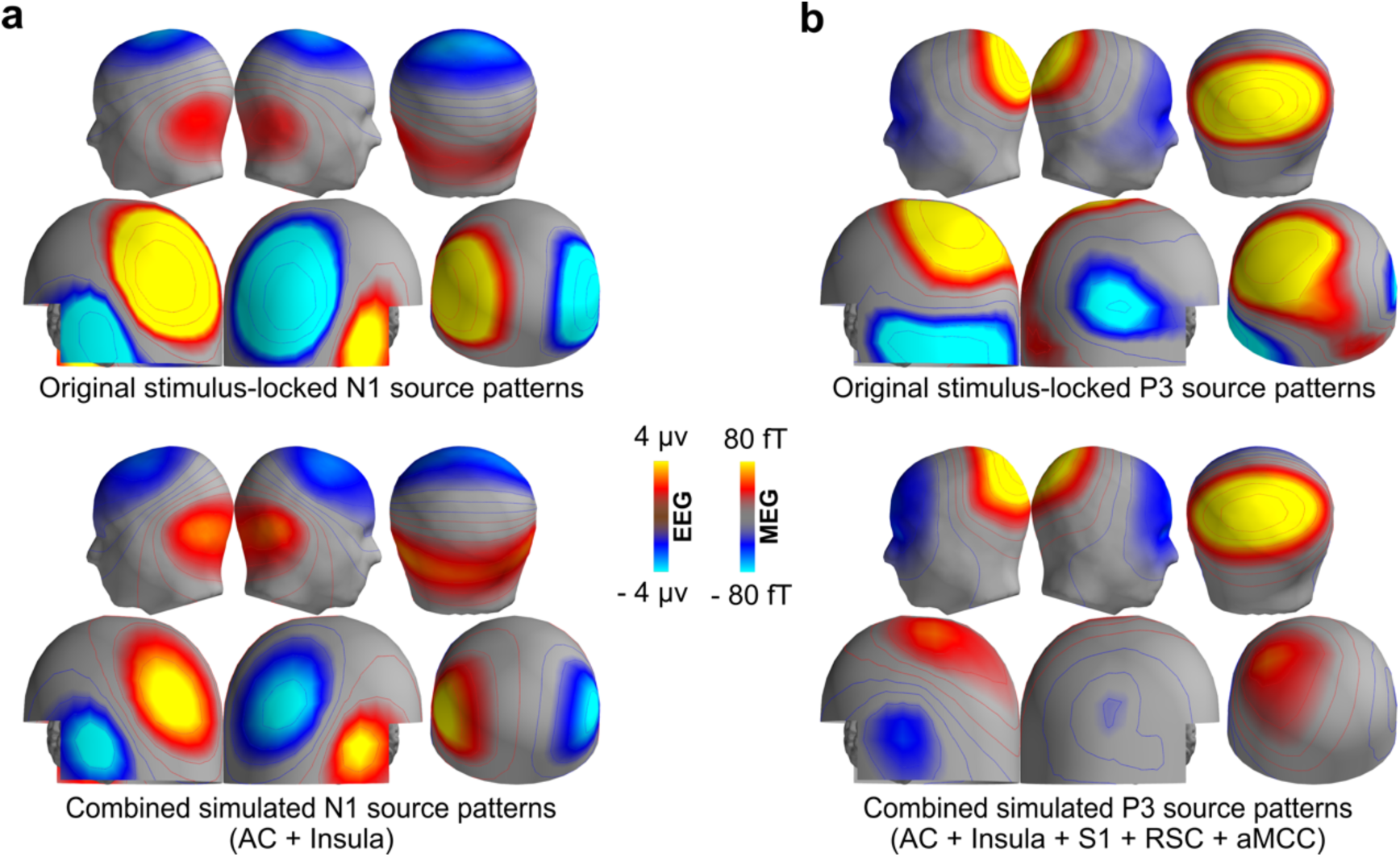
Multi-source simulation for the N1 and P3. (a) Grand average scalp EEG maps and virtual MEG magnetometer maps at the peak latency of the N1 in Cz (upper). Combined simulation of N1 maps in scalp/sensor space based on the sum of the sources in auditory cortex (AC) and insula; cf. Table S1) fitted to the stimulus-locked N1 (lower). (b) Grand average scalp EEG maps and virtual MEG magnetometer maps at the peak latency of the P3 in Pz (upper). Combined simulation of P3 maps in scalp/sensor space based on the sum of the sources in AC, insula, primary somatosensory cortex (S1), retrosplenial cortex (RSC), and anterior midcingulate cortex (aMCC) (cf. Table 1) fitted to the stimulus-locked P3 (lower).

**Table 1.** Modeling of the grand-average P3 data with simulated M/EEG based on anatomically defined source regions (cf. Fig. 4b).

| Simulated source activation strengths (nAm) |  |  |  |  | Residual Variance (%) |  |
| --- | --- | --- | --- | --- | --- | --- |
| AC | Insula | S1 | RSC | aMCC | EEG | MEG |
| 5.5 | 7.8 | 11.3 | 44.3 | 12.8 | 6.0 | 62.5 |
| - | 3.0 | 12.8 | 42.0 | 20.3 | 6.0 | 65.3 |
| 3.5 | - | 11.8 | 42.0 | 17.8 | 5.0 | 67.0 |
| 9.3 | 11.3 | - | 49.8 | 16.3 | 6.4 | 81.5 |
| 0.0 | 0.0 | 22.8 | - | 4.0 | 52.9 | 87.1 |
| 10.3 | 15.5 | 12.0 | 45.0 | - | 8.3 | 60.5 |

AC, auditory cortex; S1, primary somatosensory cortex; RSC, retro-splenial cortex; aMCC, anterior midcingulate cortex; EEG, electroencephalography; MEG, magnetoencephalography.

To test the validity of the modeling approach, the same sources were fitted to the N1 peak, leading to zero weights for all sources except for AC and insula (Fig. 5; Table S1). This model resulted in a residual variance of 16.3% in EEG and 27.8% in MEG. While the weighting of the insular cortex of about 1/3 of the auditory cortex appears relatively high, leaving out the insular ROI leads only to a minor increase of the residual variance, whereas the insular cortex alone cannot explain the N1 data well.

Finally, we tested two previous hypotheses for the generation of P3, which were not suggested by the dSPM analysis. First, a source in TPJ has been suggested based on fMRI. To this end, we used the region provided by a standard parcellation (Destrieux et al., 2010). Such a bilateral TPJ source produces a bilateral posterior maximum (Fig. S8b), but cannot replace the RSC in direct comparison. When added as the sixth source to the model from Table 1, TPJ receives no weight. When TPJ is used to replace RSC in a model with five sources, the residual variance increases to 50.4% in EEG and 85.4% in MEG, providing no support for a relevant contribution of TPJ to the P3 in the present data.

Second, a previous EEG study suggested that a distributed source in superior parietal cortex (Moores et al., 2003) could explain the P3. For this simulation, it was assumed that a distributed source existed right below the centro-posterior P3 in the EEG map, extending down to the IPS with a homogeneous amplitude distribution. This extended source indeed produces an EEG pattern with considerable similarity to the centro-posterior P3 (Fig. S8b), as well as to the RSC simulation. Replacing the RSC with this distributed source accordingly produces only slightly higher residual variance in comparison (Table S2). It is interesting to note that the dSPM estimate of this simulated source shows considerable spread to the RSC and PCC (Fig. S8a), but, conversely, no strong activity in parietal sulci was observed in the P3 source analysis, but would have been predicted by this simulation.

## 4. Discussion

Our results provide evidence of a role for RSC (Vogt, 2019; Vogt et al., 1995) in the generation of the classical P3. A second source with a typical P3 time course was observed in insular cortex, but this component was not dominant for the M/EEG maps. Other sources like S1, AC and aMCC are active in an overlapping time range but contribute to different aspects of the evoked response, which we do not consider part of the classical P3: In case of S1, the response is clearly linked to the button presses. AC and aMCC produce negative-going responses on the dorsal scalp, and their waveform and peak latencies differ from the prototypical P3 at Pz. This model is at odds with the long-held assumption that the P3 as observed by M/EEG is generated by a more distributed set of sources (Bledowski et al., 2004; Christoph Mulert et al., 2004) that is not well accessible to source analysis techniques. The results are based on the combination of EEG, MEG, and individual anatomy to provide the best possible information for the source analysis (Molins et al., 2008). Conversely, we did not directly constrain the source analysis with information from fMRI (Bledowski et al., 2004). In our view, caution is warranted when using such priors unless correlation between the brain activity measured by the different modalities has been independently confirmed; otherwise, such priors have the potential to mislead M/EEG analyses and lead to incorrect inferences regarding M/EEG sources. Indirectly, however, we used the information provided by fMRI, which first confirms that the RSC is active during target detection (Kim, 2014), and second allows for the disambiguation of the ventral RSC from the dorsal PCC, an inference that cannot be easily made based on the M/EEG data alone.

The limitations of the inverse problem remain, though, and alternative source models can easily be constructed. For example, an extended positive-going source directly below the P3 maximum in the EEG map produced a very similar map and could be used to substitute the RSC source in our model. A previous EEG study that used minimum-norm source reconstruction without noise normalization had proposed such a solution (Moores et al., 2003). However, the latter method generally prefers superficial sources (Lin et al., 2006b), which is balanced by noise normalization as used in the present study. One further difficulty of M/EEG source analysis is that, in contrast to fMRI, the distribution of a source is not directly related to the actual extent of the activity on the cortex. The pattern with opposite polarity with respect to the (outward) cortical normal in adjacent banks of a sulcus is often caused by spread of a focal source (Fig. 4b), whereas a physiological source that extends across both banks with the same polarity would lead to major signal cancelation for distant recordings in M/EEG (Goldenholz et al., 2009). We therefore chose to display source activity together with polarity information, to avoid the impression of extended sources where they are unlikely, based on the activation pattern, and estimated the spread and activation pattern of each source by simulation studies.

Generally, other sources, in particular those that contribute little or variably to either the MEG or EEG, cannot be excluded by our approach. For example, we were not able to find components of the Bereitschaftspotential and motor activity preceding the button press (Shibasaki and Hallett, 2006). Based on timing and source location, we could only identify the reafferent activity in S1 with a peak about 40 ms after the button presses (Praamstra et al., 1996), but not the motor-cortex activity directly preceding the button presses (Bötzel et al., 1993). While the response-locked source waveforms in RSC and insula have some similarity to the gradually increasing Bereitschaftspotential, different sources in SMA and motor cortex have been reported for the latter (Erdler et al., 2000; Praamstra et al., 1996). In the present study, SMA activity was observed in fMRI, but merged with more prominent activity in pre-SMA and aMCC. The time course of the related M/EEG activity was also response locked, but with an onset after the button presses and long persistence thereafter, which is not typical for the Bereitschaftspotential (Shibasaki and Hallett, 2006). The reason for not finding motor-related activity is probably that its amplitude is small relative to the P3 in the oddball paradigm.

While the source analysis was based on combined M/EEG, there are clearly differences between the two modalities. The constrained, sparse source model with five sources can achieve better residual variance in EEG than in MEG, even when considering that the residual variance of these models must not be directly compared between MEG and EEG. First, higher residual variance in MEG is generally expected based on the more focal signal in planar gradiometers, which leaves other sensors with less signal but similar noise. Second, the relative weighting of MEG and EEG is based on Z-scores, which then results in an advantage for EEG because of the higher signal to noise level. The relative amplitude scaling of MEG and EEG simulations depends on assumptions made in the head model, in particular the conductivity of the EEG model, which is individually different and difficult to estimate exactly. As a consequence, the amplitude of the MEG is somewhat underestimated by the model, for both N1 and P3, which is an additional source of higher residual variance in MEG compared to EEG. Residual variance in MEG was accordingly higher for the N1 model, as well, but more prominently for the P3 model. This is partly related to the AC source of the N1 producing a tangential source, with excellent detection in MEG as well as EEG. In contrast, the RSC source, which explains most of the variance in the P3 model, has a predominantly radial orientation, which produces comparatively weak MEG signals. As a consequence, the MEG maps were more variable and often dominated by the source in S1. Our highly constrained model cannot explain the full variance of the MEG, which is partly explained by the points listed above, but the potential need for additional sources that are missing in the model used in this paper cannot be excluded. For example, we observed some activity in V1 and the POS in M/EEG and fMRI, which we did not include into the model. The context of this occipital activity remains unclear at this point; V1 activity could e.g., be related to response-locked blinks, but in the case of M/EEG could as well be spread from RSC (cf. Fig. 4a).

The other limitation is the degree to which fMRI can be used to constrain M/EEG source analysis. All sources that were found active in the dSPM maps of this study were also confirmed by fMRI (the potential exception is discussed later), but fMRI shows activity in additional areas that were not revealed by M/EEG source analysis. There are at least two potential sources for this discrepancy: First, electric activity of similar surface polarity cancels out for M/EEG if the source spans two sides of a sulcus (Ahlfors et al., 2010), whereas this configuration would rather support the activity’s detection in fMRI. This could e.g., apply to potential sources in the TPJ; it could then be that there was another P3 source in this region, but that its signal-to-noise ratio was too low or its source location too variable to be detected by cross-subject M/EEG. Second, fMRI often does not match with low-frequency M/EEG activity in the delta and theta band such as e.g. the error-related negativity (Agam et al., 2011), but better with neural activity in the gamma band (Logothetis et al., 2001; Niessing et al., 2005; Steinmann and Gutschalk, 2011). Thus, while the strong RSC activity in fMRI generally supports the RSC’s contribution to the P3 source, it still remains possible that the relationship between M/EEG and fMRI activity in this region is indirect, e.g., via functionally coupled gamma activity.

Activity in the PCC and RSC has also been reported in intracranial recordings of the P3 (Halgren et al., 1995b), without providing a clear separation between the two. While the authors of that study suggested that the PCC/RSC was a source of the P3a rather than the P3b, neither fMRI (Kim, 2014) nor source-analysis studies (Bledowski et al., 2004) have confirmed such a strict separation of P3 subcomponents as suggested based on these intracranial data. As a limitation, while the depth recordings found high amplitudes in and near PCC, no polarity reversal was found (Halgren et al., 1995b), which would have confirmed that the electrode passed through the source.

An additional case for a potential P3 source in RSC is how this region is connected to other brain networks recruited by target processing. The PCC has been demonstrated to be a major hub of the “task-negative” default-mode network (Fox et al., 2005), while activation during oddball detection (Kim, 2014) has been observed in “task-positive” networks (Fox et al., 2005; Hugdahl et al., 2015) such as the dorsal and ventral attention networks (Yeo et al., 2011). In the early resting-state network studies, PCC activation included all of RSC (Fox et al., 2005), whereas later, more detailed network maps (Yeo et al., 2011) segregated the dorsal part of RSC into a fronto-parietal network, which would better match with a role in active target detection. Anatomical studies in monkeys indicate that both PCC and RSC are reciprocally connected with multiple frontoparietal areas that are active during oddball tasks in fMRI (Kobayashi and Amaral, 2007; Vogt and Pandya, 1987). This would be consistent with the idea that even if the fronto-parietal network does not itself generate the P3, it may still be functionally coupled with a generator in RSC. Such connectivity would explain previous findings of reduced P3 with right-TPJ lesions (Knight et al., 1989; Verleger et al., 1994), even if TPJ was not the source of P3. In fact, a model where TPJ provides input into RSC, the neuroelectric source of P3, could better explain why unilateral TPJ lesions caused bilateral reduction of the P3 (Knight et al., 1989).

The RSC is also functionally coupled to the hippocampus in the medial temporal lobe (Alexander et al., 2018). Given the hippocampal P3-like activity demonstrated by iEEG (Halgren et al., 1980), this raises the possibility of a close functional coupling between the extracranial P3 in M/EEG and the intracranial hippocampal activity, despite their anatomical dissociation. Another important question for the source analysis and simulation studies was if hippocampal P3-like activity could potentially be recorded in M/EEG. Despite its clear demonstration in iEEG (Halgren et al., 1980), no hippocampal activity has been shown in fMRI in this (Fig. 2c) or previous odd-ball-paradigm fMRI studies (Kim, 2014). One possible reason for this negative finding could be different neuro-vascular coupling in medial temporal lobe compared to neocortex (Hill et al., 2021), suggested recently based on combined iEEG and fMRI. While the contribution of a hippocampal source to the parietal P3 in EEG had already been excluded based on lesion studies (Johnson, 1988; Onofrj et al., 1992), this does not exclude that hippocampal activity may generally contribute to other aspects of the M/EEG response (Alberto et al., 2021), even though signal-to-noise ratios for such areas are weak. Indeed, the mapping shown in Fig. 2 also suggests activity in the medial temporal lobe. However, this activity was as prominent in the N1 as in the P3 time interval, which is not consistent with known iEEG time courses in hippocampus (Halgren et al., 1980). Moreover, we demonstrated that there is considerable spread and crosstalk between the medial temporal lobe and AC as well as insular cortex (Fig. 4). It is therefore more likely that the activity observed in the medial temporal lobe in our source analysis represents spread from AC and insular cortex, particularly given the fact that M/EEG signal-to-noise ratio is much higher in AC (and somewhat higher in insular cortex) than in the hippocampus (Goldenholz et al., 2009). This leaves us with the paradoxical situation that there is strong iEEG evidence for P3-like activity in the hippocampus evoked by the paradigm used (Halgren et al., 1980), but that this activity is hard to detect or to distinguish from other sources with all three non-invasive techniques used in this study.

The situation is somewhat different for the insula. While there is also spread from AC to the insula (or its vicinity) in the N1 time interval, the pattern is clearly different in the P3 time interval, with surface-positive activity in the insular cortex; activity in AC remains surface negative in this time interval, as reported previously for a passive oddball paradigm (Kretzschmar and Gutschalk, 2010). We therefore consider it more likely that the P3-like time course shown in Fig. 3 is generated in the insula, rather than in the temporal lobe. P3 generators in the insula have been suggested before. An EEG study (Bledowski et al., 2004) suggested a contribution of insular cortex to the P3a. A recent iEEG study demonstrated a stronger P3b in anterior insular cortex (Citherlet et al., 2020), in synchrony with gamma activity in the same latency range. Given the observation of strong fMRI activity for detected oddballs in insular cortex, this supports the hypothesis stated above that gamma activity is a potential link between the P3 and BOLD activity. Finally, strong fMRI activity was observed in aMCC. Insular and aMCC activity show high functional connectivity in fMRI (Yeo et al., 2011), but the time course of the insula and aMCC found here are quite different: the insular time course is similar to the RSC and shows a build-up towards the time of the button press. In contrast, aMCC activity was most prominent after the button press and may thus indicate some kind of performance control (Heilbronner and Platt, 2013).

At this point, this source analysis is limited to a single paradigm, the classical auditory oddball paradigm. The dominant component in this paradigm is the P3b, but it can be expected that some P3a source activity will also be present. Therefore, we cannot as yet make strong conclusions with respect to the neural sources of these subcomponents. The EEG distribution of the simulated bilateral RSC source over centro-parietal electrodes, however, makes it a better fit for the P3b rather than the P3a (Polich, 2007). Dissociating if other sources are more specific for the P3a (Halgren et al., 1995a) or if there is strong overlap in the generators of these two subcomponents (Kim, 2014) will require further studies that manipulate the relative strength of these components. Other, more complex tasks will certainly be expected to involve additional brain regions. While we propose that the P3 generator in RSC will remain a constant contributor for such paradigms as well, this hypothesis requires evaluation in future experiments or the reevaluation of existing data.

## 5. Conclusion

Multiple neural processes are active in parallel with the P3 in M/EEG, some observed more easily with fMRI and some more easily with EEG or MEG. But while the P3 is most likely functionally coupled to this distributed neural network, it does not appear to be the bioelectric source of the classical, parietal P3 signal measured in EEG. Based on the evidence presented here, this source appears to be more focal and to lie in the RSC. This finding is essential to explore the functional role of the P3 between the fronto-parietal network observed in fMRI (Kim, 2014) and the hippocampal P3-like activity demonstrated with iEEG (Halgren et al., 1980), and will help to better understand the functional roles of both RSC and the P3. Moreover, understanding its functional anatomy may support the application of the P3 as diagnostic tool. For example, reduced P3 in Alzheimer’s disease (Frodl et al., 2002) might be linked to cortical hypometabolism and tau accumulation (Strom et al., 2022), the latter of which has been suggested to covary with the connectivity between RSC and hippocampus (Ziontz et al., 2021). We hope that future invasive studies will seek to confirm the source configuration suggested by this non-invasive study, possibly by demonstrating co-occurrence of (high-)gamma activity together with a typical P3 time course in RSC.

## Acknowledgements

This work was primarily supported by Deutsche Forschungsgemeinschaft grant DFG 593/5-1 (AG) and Bundesministerium für Bildung und Forschung grant 01EV0712 (AG), as well as by National Institutes of Health grants R01NS104585 and P41EB030006 (MSH).

## Abbreviations

AC: auditory cortex
aMCC: anterior midcingulate cortex
IPS: intraparietal sulcus
MCC: midcingulate cortex
PCC: posterior cingulate cortex
POS: parieto-occipital sulcus
ROI: region of interest
RSC: retro-splenial cortex
S1: primary somatosensory cortex
SMA: supplementary motor area
TPJ: temporo-parietal junction;
V1: primary visual cortex.

## Author contributions

**Diptyajit Das:** Conceptualization, Formal Analysis, Data Curation, Visualization, Writing - Original Draft; **Marnie E. Shaw:** Conceptualization, Investigation, Writing - Review & Editing; **Matti S. Hämäläinen:** Validation, Writing - Review & Editing; **Andrew R. Dykstra:** Validation, Writing - Review & Editing; **Laura Doll:** Validation, Writing - Review & Editing; **Alexander Gutschalk:** Conceptualization, Supervision, Validation, Writing - Original Draft & Editing, Funding acquisition.

## Data availability

The raw and processed M/EEG and fMRI data and scripts will be made available on heiDATA, the open research data repository of Heidelberg University under the following doi: https://doi.org/10.11588/data/YB9SQI

**Appendix A.** Supplementary material

