## Appendix for "A role for retro-splenial cortex in the task-related P3 network"

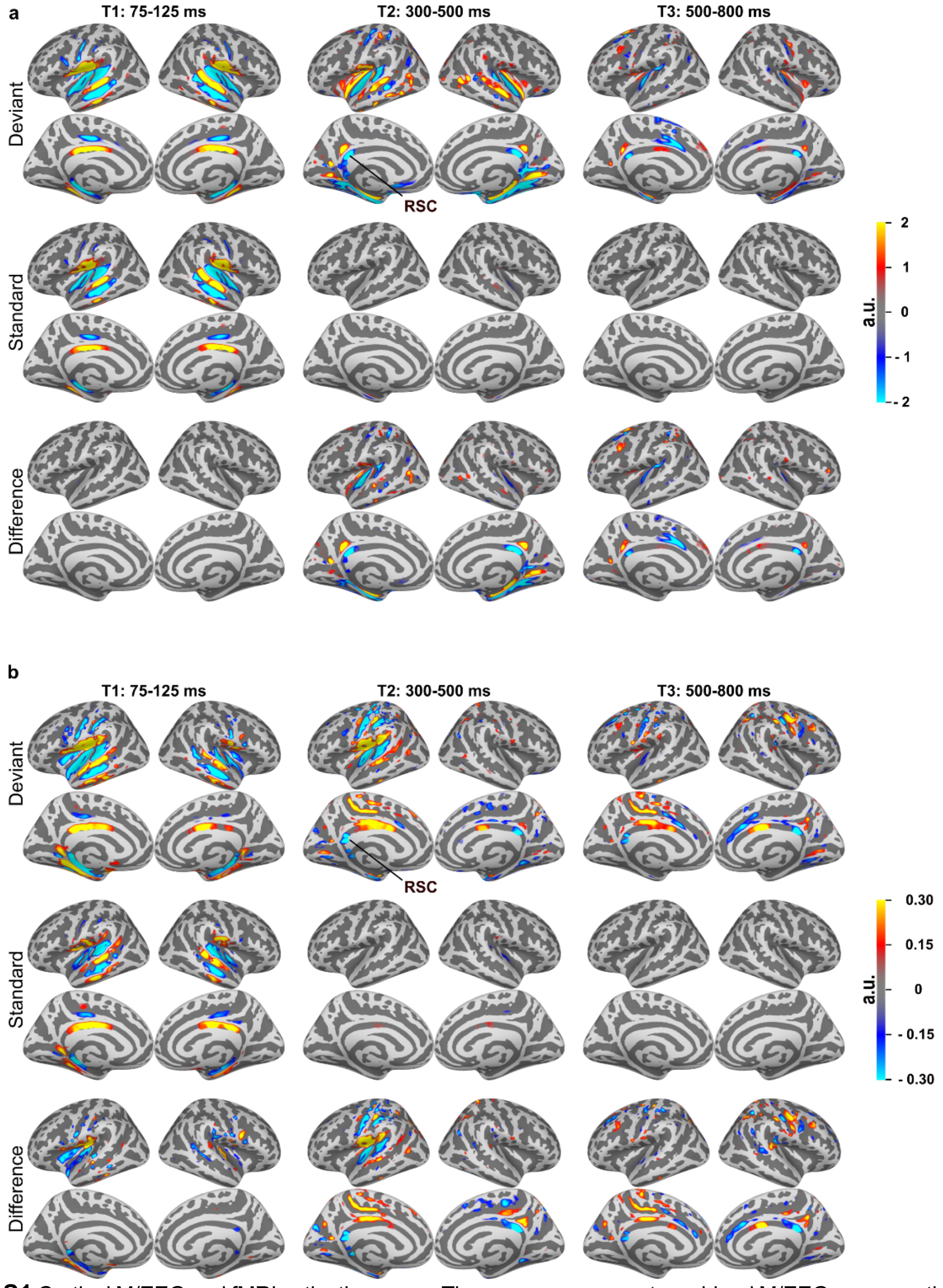


**Fig. S1** Cortical M/EEG and fMRI activation maps. These maps represent combined M/EEG source estimates similar to those shown in Figure 2a, but with alternative source analysis methods, using (a) sLORETA (Pascual-Marqui R D, 2002) and (b) a Borgiotti-Kaplan beamformer (Sekihara and Nagarajan, 2008). Note that the numerical values cannot be directly compared between these source estimation methods, even though they represent noise normalized z-scores.

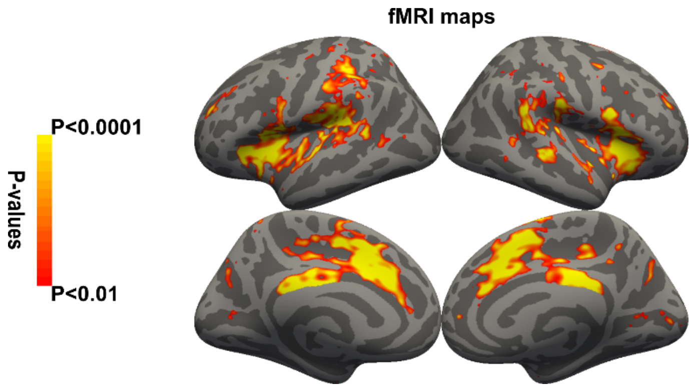


**Fig. S2** fMRI maps for the contrast deviants – standards, based on a random-effects statistic. Same analysis as Figure 2c but with more conservative cutoff (p<0.01, FDR corrected).

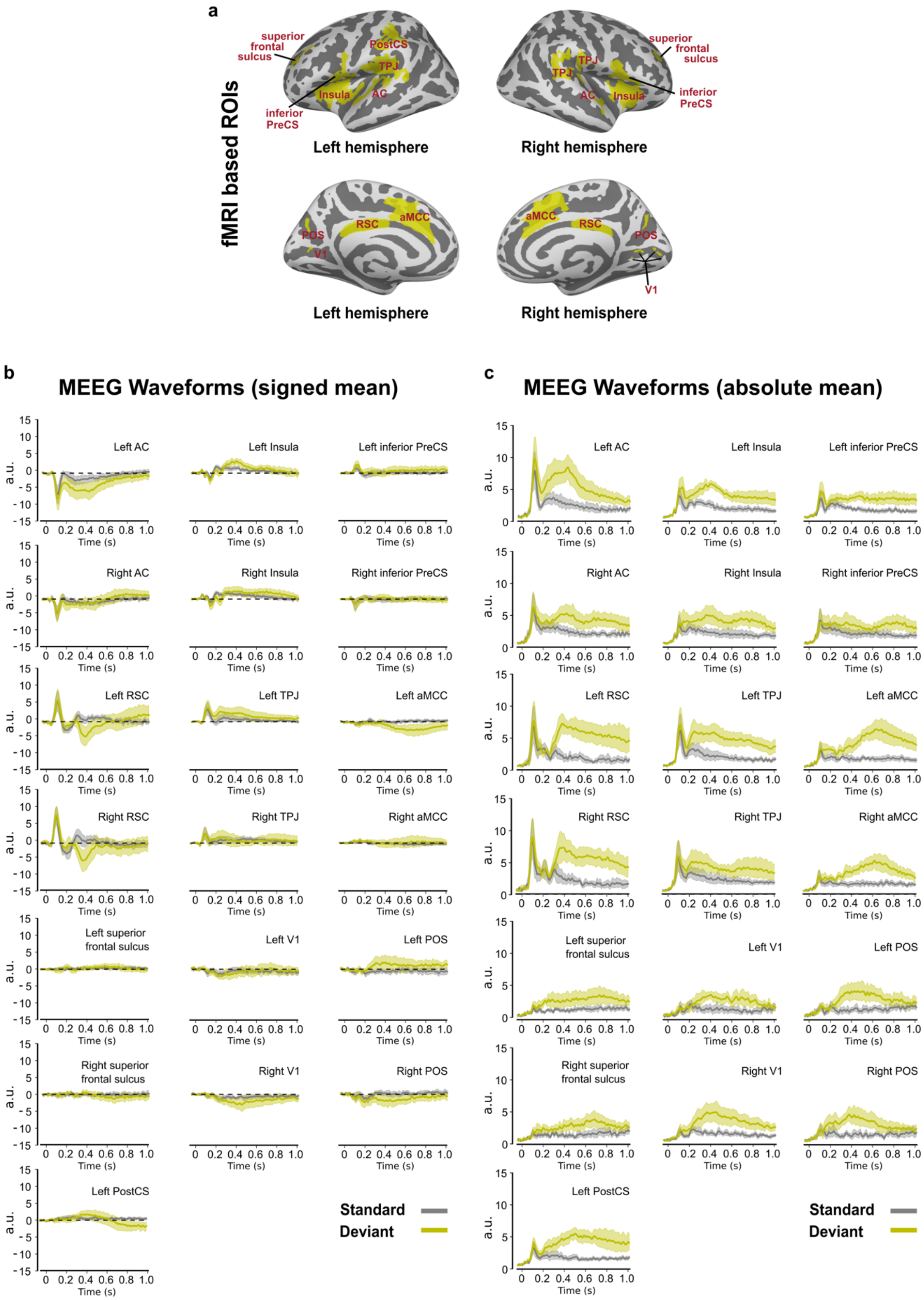
a
fMRI based ROIs
superior frontal sulcus
PostCS
TPJ
Insula
AC
inferior PreCS
Left hemisphere
Right hemisphere
aMCC
RSC
POS
V1
b
MEEG Waveforms (signed mean)
c
MEEG Waveforms (absolute mean)
Left AC
Left Insula
Left inferior PreCS
Right AC
Right Insula
Right inferior PreCS
Left RSC
Left TPJ
Left aMCC
Right RSC
Right TPJ
Right aMCC
Left superior frontal sulcus
Left V1
Left POS
Right superior frontal sulcus
Right V1
Right POS
Left PostCS
a.u.
Time (s)
Standard
Deviant

**Fig. S3** M/EEG time course analysis with ROIs based on the cortical fMRI activation. The ROIs are based on a cutoff of $p<0.01$ (cf. Fig. S2). (a) Connected activation clusters were separated to restrict each pattern to a defined cortical region. The time courses in these ROIs were calculated in two different ways: (b) as signed waveform, where the polarity of each time point is maintained. To avoid signal cancelation, the orientation of dipoles that deviate from the average orientation of the ROI are inverted with the "mean flip" procedure in MNE python. Upward going waveforms have an overall positive polarity at the cortical surface, downward going waveforms have a negative polarity at the cortical surface (c) the mean is calculated for the absolute values. In both cases, the anatomical ROIs are defined based on the average fMRI maps, but are then transformed back to the individual anatomy. Here, the waveforms are extracted and averaged within the individual anatomical selection of dipole sources.

The methods in (b) and (c) have different advantages and problems. (b) Preserves the polarity information and is less susceptible for spread from other areas. It may overlook activity with a different orientation, which could be a problem if the ROI was designed with the wrong orientation. It is also less sensitive for cases where the inter-individual variability of the source location, orientation, or cancelation may be high (c) is very sensitive for the latter cases instead, but is also more susceptible to spread from other areas. There is generally an enhancement of noise. Waveforms appear changed, because e.g. subsequent negative- and positive-going parts of the waveforms are added, producing waveforms that are hard to recognize in the original, scalp and sensor waveforms. A short comment and interpretation considering the weaknesses and strengths of the two methods is provided below for all ROIs:

Auditory cortex (AC): Typical N1 followed by a negative-going sustained field that is stronger on the left. The sustained field looks more "peaked" and less "sustained" in the absolute waveforms but the two are otherwise similar.

Retrosplenial cortex (RSC): Negative-going peak around 400-ms latency that is only observed for deviants. In the absolute-mean, the onset of the peak shows a similar waveform and latency, but it declines only slowly and shows a very sustained waveshape. This could e.g. be related to spread from the longer-latency activity in more anterior parts of cingulate cortex. As in many other ROIs, strong activity around 100-ms latency is observed, which is supposedly spread from auditory cortex. Overall, activity in this fMRI-based ROI is similar to the M/EEG based ROI and further supports a P3 generator in RSC. Note that the waveform is negative going, but that the orientation of the cortical surface in RSC is oriented towards the ventral side of the brain.

Superior frontal sulcus: Almost no activity in the signed waveforms; the absolute waveforms show a sustained shift for deviants over standards and maybe a weak peak after 600 ms. This activity could possibly be related to the stronger activity seen in aMCC, but does not reflect the waveform of the scalp P3.

Left post-central sulcus (PostCS): Activity in this ROI is very likely related to somatosensory activity. This source is strictly left lateralized, in both, fMRI and MEG.

Insula: A P3-like waveshape is observed in both, signed and unsigned waveforms, but the peak is comparatively weak. Overall, the waveform appears similar to the waveform in the M/EEG based ROI.

Temporo-parietal junction (TPJ): Positive-going activity with an early peak after 100 ms, followed by a sustained wave, which is stronger for deviants. Overall, this waveform resembles activity from auditory cortex. Based on the polarity and the ventral orientation of the ROI, the activity in the middle of the scalp would be expected to be negative going. Accordingly, we think that this ROI mostly reflects spread from the auditory cortex, and is unlikely to be a major generator of the scalp P3.

Primary visual cortex (V1): This ROI reflects some negative-going activity in the P3 time range, somewhat stronger on the right and similar in signed and absolute waveforms. A neural generator contributing to the P3 in this area is therefore possible, but the physiological context to the P3 remains unclear. Possible explanations could be a relationship to the task response and correlated blinks or eye movements, which might evoke activity in fMRI as well as M/EEG. However, a direct relationship to the P3 network cannot be excluded (even if we consider it unlikely).

Pre-central sulcus (PreCS): This activity is very close to the activity cluster in insula in fMRI. Nevertheless, the waveforms in this ROI do not show any P3-like waveforms in the signed analysis. Even in the absolute waveforms, there is more of a sustained shift, and only a subtle P3-like wave on top, which is much weaker than in the neighboring insula.

Anterior midcingulate cortex (aMCC): In contrast to the M/EEG based ROI, this fMRI-based ROI also includes the pre-supplementary motor area (pre-SMA). The waveforms are nevertheless similar to the M/EEG ROI, with a negative-going, long-latency wave that is stronger on the left. This lateralization is not so clear in the absolute-value analysis, but otherwise the waveforms are similar.

Parieto-occipital sulcus (POS): A positive-going wave is seen on the left and a negative going wave on the right, both with a P3-like waveform and latency. This finding matches well with the dSPM for the time range 300 - 500 ms, which shows positive-going activity in left POS. The right-hemisphere waveform may therefore reflect spread from the close-by left-hemisphere. Alternatively, the orientation of the activity may be very variable across participants, e.g. because the generator is located on both sides of the sulcus, producing very different patterns between participants depending on individual anatomy and resulting patterns of cancelation. At this point, these different explanations cannot be resolved. It is therefore possible that another P3 generator exists in POS, but the reason for the left-dominance observed in this sample would still remain unclear.

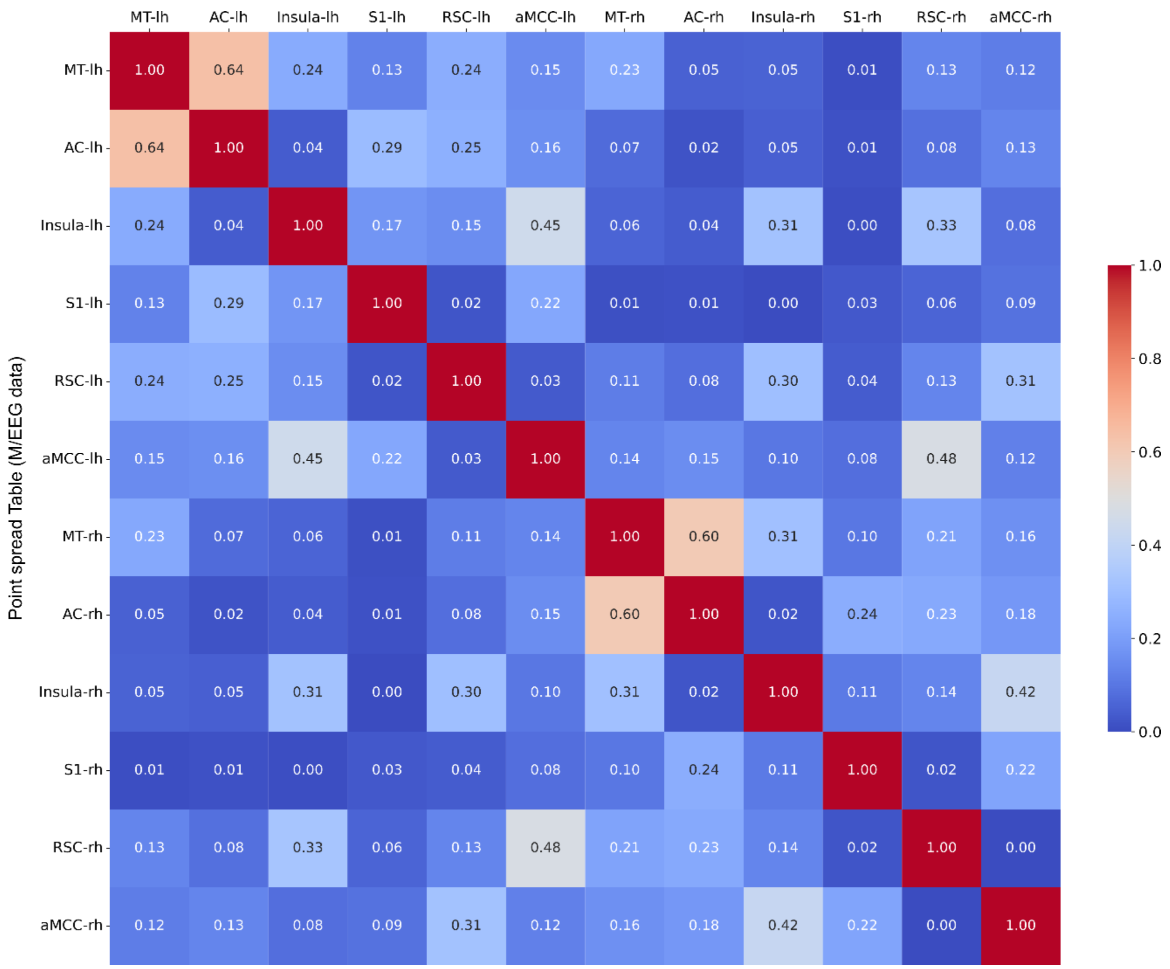


**Fig. S4** Point spread analysis table based on the M/EEG data. The spread values range from 0 (minimum) to 1 (maximum).

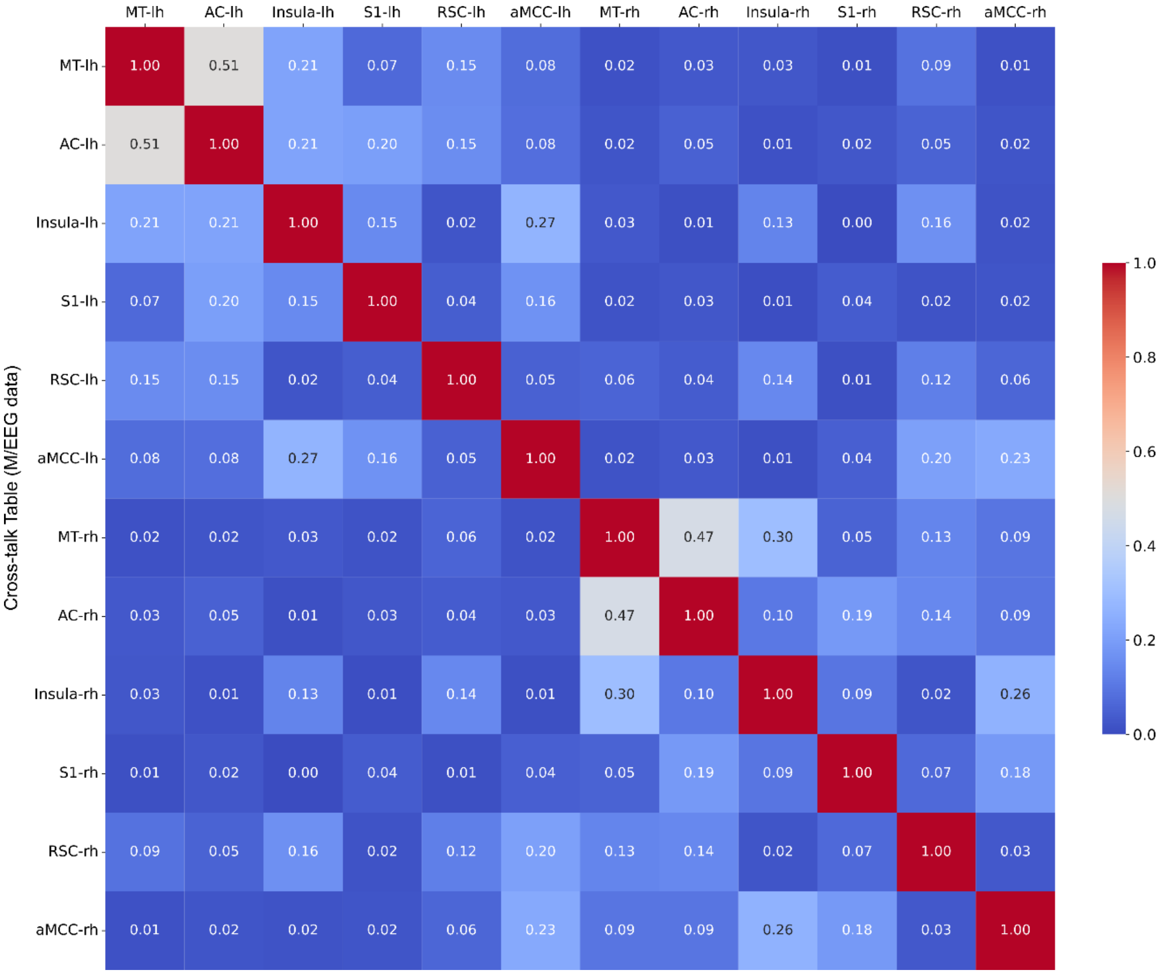


**Fig. S5** Cross-talk analysis table based on the M/EEG data. The cross-talk values range from 0 (minimum) to 1 (maximum).

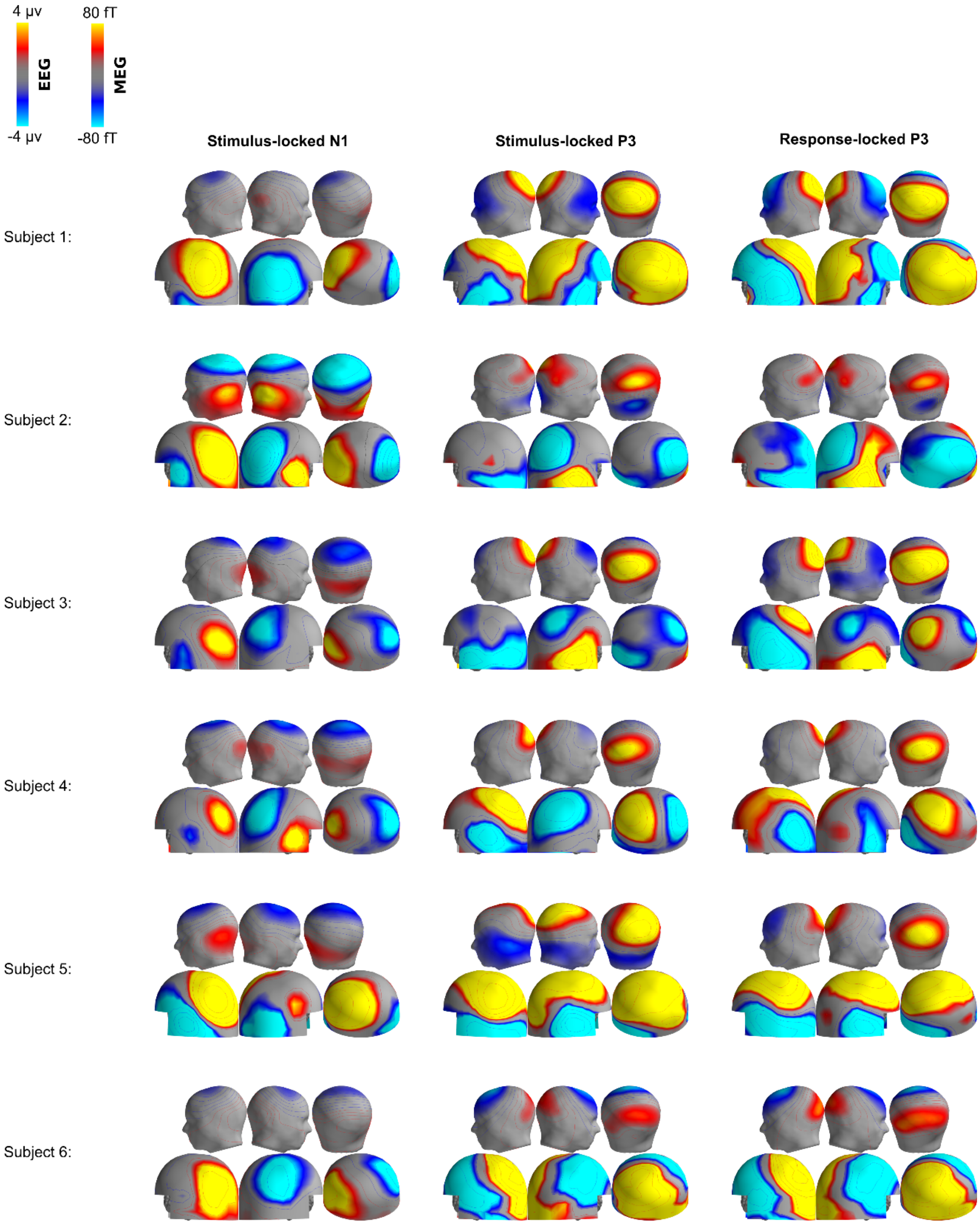


**Fig. S6** EEG and MEG maps for individual participants. Sensor-level activities are mapped at the individual peak latency for N1 (left) stimulus-locked P3 (middle), and response-locked P3 (right) for individual participants 1 - 6.

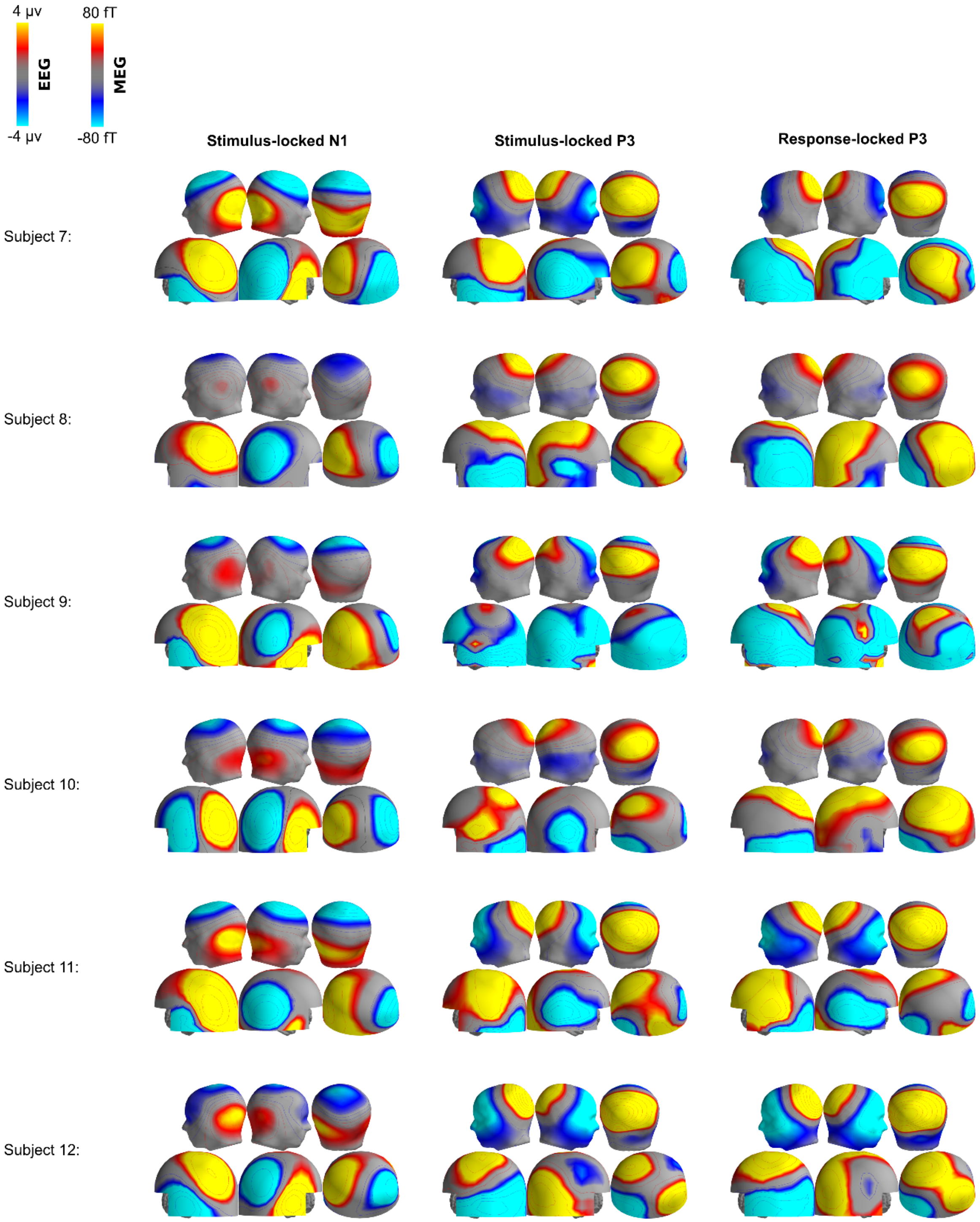


**Fig. S7** EEG and MEG maps for individual participants. Sensor-level activities are mapped at the individual peak latency for N1 (left) stimulus-locked P3 (middle), and response-locked P3 (right) for individual participants 7 - 12.

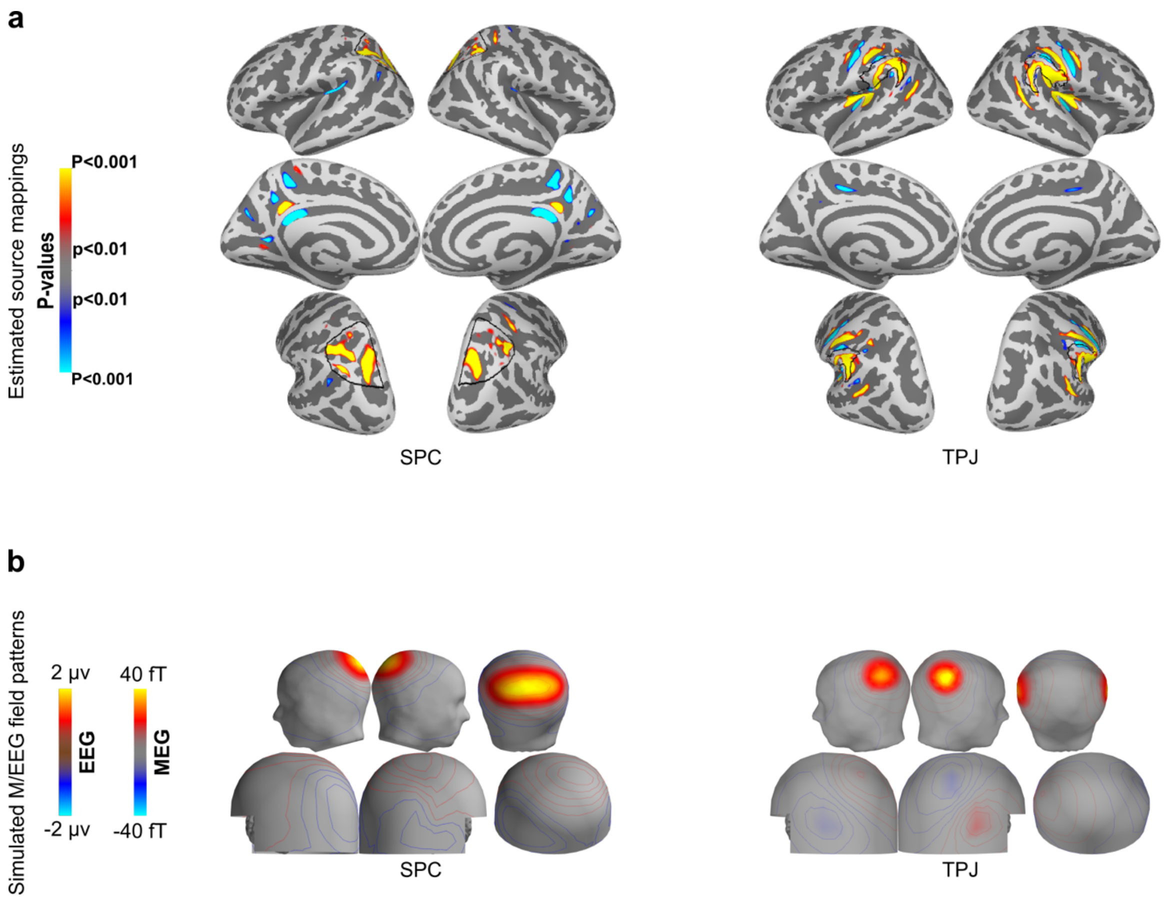


**Fig. S8** Simulated M/EEG for a distributed superior parietal cortex (SPC) and a temporo-parietal junction (TPJ) source. (a) dSPM analysis for simulated distributed SPC (left) and TPJ (right) sources (n=12 subjects; p<0.01). (b) Average EEG and MEG maps for the simulated distributed SPC (left) and TPJ (right) sources.

**Table S1**

Modeling the grand-average N1 pattern with simulated M/EEG data based on anatomically defined source regions (Figure 4b).

| **Simulated source activation strengths (nAm)** | | | | | **Residual Variance (%)** | |
|---|---|---|---|---|---|---|
| **AC** | **Insula** | **S1** | **RSC** | **aMCC** | **EEG** | **MEG** |
| 27.8 | 9.3 | 0.0 | 0.0 | 0.0 | 16.3 | 27.8 |
| 28.8 | - | - | - | - | 20.7 | 26.8 |
| - | 26.5 | - | - | - | 69.6 | 100.0 |

AC, auditory cortex; S1, primary somatosensory cortex; RSC, retro-splenial cortex; aMCC, anterior midcingulate cortex; EEG, electroencephalography; MEG, magnetoencephalography.

**Table S2**

Modeling the grand-average P3 pattern with simulated M/EEG data based on anatomically defined source regions (Figure 4b). In this simulation, the extended superior parietal source (SPC; see Figure S7) is used instead of the retro-splenial cortex (RSC).

| Simulated source activation strengths (nAm) | | | | | Residual Variance (%) | |
|---|---|---|---|---|---|---|
| **AC** | **Insula** | **S1** | **SPC** | **aMCC** | **EEG** | **MEG** |
| 5.0 | 11.3 | 10.0 | 69.5 | 0.0 | 8.9 | 66.6 |
| - | 7.3 | 11.8 | 66.3 | 6.5 | 8.8 | 70.3 |
| 3.0 | - | 11.3 | 64.0 | 6.3 | 8.0 | 73.4 |
| 9.0 | 16.0 | - | 79.0 | 0.0 | 11.1 | 75.1 |
| 0.0 | 0.0 | 22.8 | - | 4.0 | 52.9 | 87.1 |
| 5.0 | 11.3 | 10.0 | 69.5 | - | 8.9 | 66.6 |
| 5.0 | 11.3 | 10.0 | 69.5 | 0.0 | 8.9 | 66.6 |

AC, auditory cortex; S1, primary somatosensory cortex; RSC, retro-splenial cortex; aMCC, anterior midcingulate cortex; EEG, electroencephalography; MEG, magnetoencephalography.
